# Systematic characterization of the local evolutionary space available to human PKR and vaccinia virus K3

**DOI:** 10.1101/2023.11.21.568178

**Authors:** Michael J. Chambers, Sophia Scobell, Meru J. Sadhu

**Affiliations:** Center for Genomics and Data Science Research, National Human Genome Research Institute, National Institutes of Health, Bethesda, Maryland, USA; Department of Microbiology & Immunology, Georgetown University, Washington DC, USA

## Abstract

The interfaces between host and viral proteins can be dynamic spaces in which genetic variants are continually pursued, giving rise to evolutionary arms races. One such scenario is found between the mammalian innate immunity protein PKR (protein kinase R) and the poxvirus antagonist K3. Once activated, PKR phosphorylates the natural substrate eIF2α, which halts protein synthesis within the cell and prevents viral replication. K3 acts as a pseudosubstrate antagonist against PKR by directly antagonizing this halt in protein synthesis, enabling poxviruses to replicate in the cell. Exploring the impact of genetic variants in both PKR and K3 is necessary not only to highlight residues of evolutionary constraint and opportunity but also to elucidate the mechanism by which human PKR is able to subvert a rapidly evolving viral antagonist. To systematically explore this dynamic interface, we have generated a combinatorial library of PKR and K3 missense variants to be co-expressed and characterized in a high-throughput yeast selection assay. This assay allows us to characterize hundreds of thousands of unique PKR-K3 genetic combinations in a single pooled experiment. Our results highlight specific missense variants available to PKR that subvert the K3 antagonist. We find that improved PKR variants are readily available at sites under positive selection, with limited opportunity at sites interfacing with K3 and eIF2α. Additionally, we find many variants that improve and disable K3 antagonism, suggesting a pliable interface. We reason that this approach can be leveraged to explore the evolutionary plasticity of many other host-virus interfaces.

## Introduction

Molecular interfaces between interacting host and pathogen proteins are atomic battlegrounds that give rise to evolutionary arms races (Daugherty & Malik, 2012). Both proteins pursue genetic variants to gain an advantage, manifesting as sequence diversity under positive selection. This phenomenon is underpinned by the ‘Red Queen Hypothesis’, stating that constant adaptation between antagonistic interacting proteins is necessary to sustain their respective functions (Van Valen, 1973). Here we investigate an ongoing evolutionary arms race between a component of our immune system, Protein Kinase R (PKR), and a vaccinia virus (VACV) pseudo-substrate antagonist, K3, both of which are under positive selection (Elde et al., 2009; Jacquet et al., 2022; Rothenburg et al., 2009). Prior studies have characterized individual residue variants in both proteins that alter their interaction (Dar & Sicheri, 2002; Kawagishi-Kobayashi et al., 1997; Seo et al., 2008). By systematically characterizing nonsynonymous variants we gain a general understanding of their evolutionary plasticity as well as a clear map of the functional and deleterious variants that are readily available.

PKR is a critical component of the human innate immune system (Balachandran et al., 2000; Clemens, 1997; Metz & Esteban, 1972; Williams, 1997). Expressed from the gene *EIF2AK2* (eukaryotic translation initiation factor 2 alpha kinase 2, also referred to as *PKR*), it is composed of two N-terminal double-stranded RNA (dsRNA) binding domains and a C-terminal kinase domain, and the two are connected by an extended linker region (Figure 1A) (Dar et al., 2005). PKR is one of four kinases that activates the Integrated Stress Response (ISR), ultimately inhibiting translation initiation to halt protein synthesis. Each kinase is activated by a distinct stress signal and converge by phosphorylating the same substrate, eIF2α (eukaryotic translation initiation factor 2 subunit alpha) (Chen & London, 1995; Dever et al., 1992; Galabru & Hovanessian, 1987; Hovanessian, 1989; Shi et al., 1998). PKR is constitutively expressed across all tissue types (GTEx Analysis Release V8, dbGaP Accession phs000424.v8.p2) but is not activated until it binds dsRNA, a general byproduct of viral replication (Roberts et al., 1976; Weber et al., 2006). Upon dsRNA binding, PKR forms a homodimer, self-activates, then phosphorylates eIF2α. Phosphorylated eIF2α inhibits eIF2B, preventing the exchange of eIF2 GDP to GTP and disrupting translation initiation (Hershey, 1991; Kaufman, 2000; Sudhakar et al., 2000). Thus, activated PKR curtails protein synthesis and

**Figure 1.**
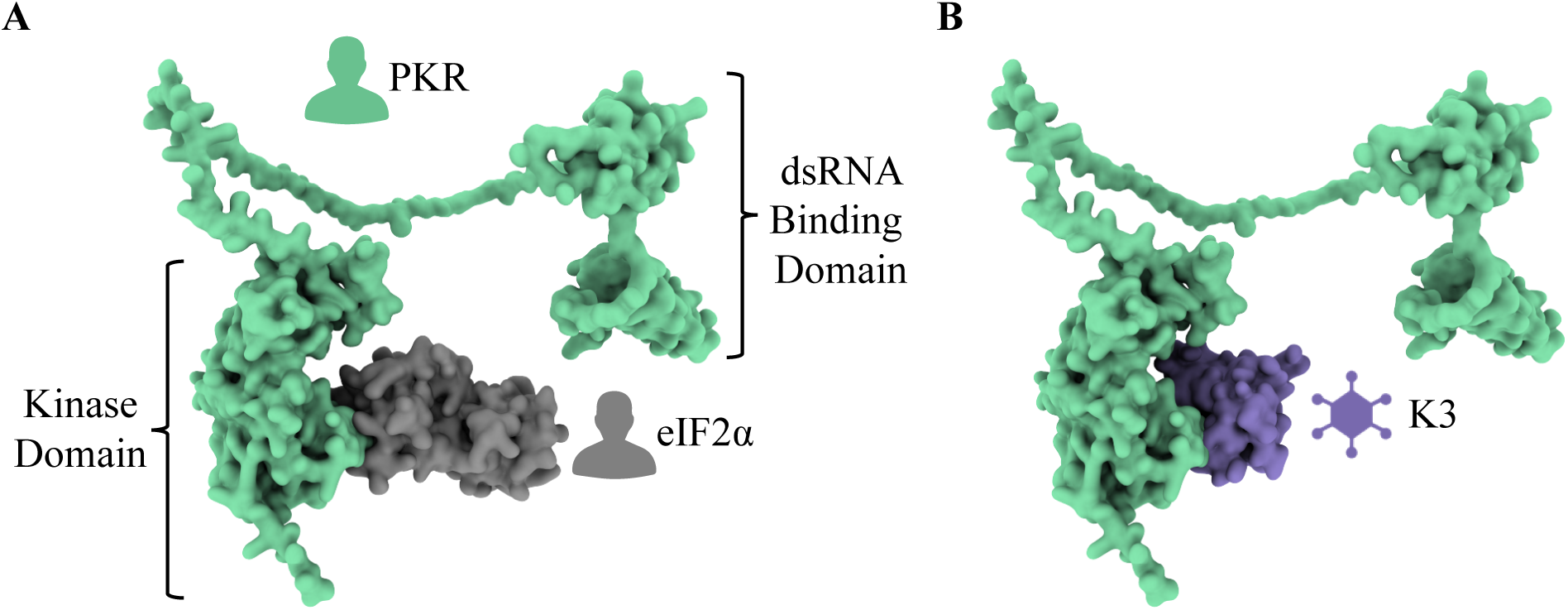
K3 acts as a pseudosubstrate antagonist by mimicking the natural substrate of PKR, eIF2α. **(A)** The human protein PKR (green) is composed of two N-terminal dsRNA binding domains and a C-terminal kinase domain. This depiction illustrates the interaction between PKR and its natural substrate, eIF2α (gray). This depiction of PKR in complex with eIF2α was generated by aligning PDB 2A1A Chain B to AlphaFold2 P19525 (see Methods). **(B)** K3 acts as a pseudosubstrate inhibitor of PKR by mimicking the natural substrate and preventing the phosphorylation of eIF2α. The predicted PKR-K3 complex was generated by aligning PDB 1LUZ Chain A to PDB 2A1A Chain A using PyMol (see Methods).

VACV K3, encoded by *K3L*, is a pseudosubstrate antagonist that is expressed by many poxviruses (Beattie et al., 1991; Haller et al., 2014). Poxviruses are large, dsDNA viruses that can infect most vertebrates and insects (Harrison et al., 2004; Moss, 2007). They replicate in the cytoplasm and generate dsRNA in the process (Moss, 1974). To prevent PKR activation by dsRNA, most known poxviruses encode proteins whose dedicated function is to inhibit PKR. VACV K3, encoded by the gene K3L, is a PKR antagonist with homologs in many poxvirus genomes (Beattie et al., 1991; Haller et al., 2014). K3 mimics the natural substrate eIF2α and acts as a pseudosubstrate inhibitor of PKR (Figure 1B) (Beattie et al., 1991; Carroll et al., 1993; Davies et al., 1993). Direct binding of K3 to PKR in place of eIF2α prevents eIF2α phosphorylation and aids viral replication by masking the virus from host PKR (Beattie et al., 1995; Elde et al., 2009).

Previous work has found both PKR and K3 to be under positive selection, with specific residues highlighted throughout PKR that are rapidly evolving (Elde et al., 2009; Jacquet et al., 2022; Rothenburg et al., 2009). Others have identified single-residue variants of PKR and K3 that alter their interaction (Kawagishi-Kobayashi et al., 1997; Seo et al., 2008). For example, the variant K3-H47R is an improved antagonist of human PKR (Kawagishi-Kobayashi et al., 1997). These findings suggest an evolutionary arms race in which improved genetic variants are continually explored (Daugherty & Malik, 2012).

We adopted a systematic approach to characterize the evolutionary landscape available to human PKR and VACV K3. We made hundreds of select, single-residue, nonsynonymous variants of *PKR* and *K3L* that screened in a high-throughput yeast growth assay. In taking this approach we were able to characterize many previously identified variants of *PKR* and *K3L* as well as several newly identified variants. We also highlight several residues under positive selection that were found to harbor functional variants of *PKR*. Our work offers some insight towards the evolutionary potential of *PKR* and *K3L* as well as a novel, systematic, and unbiased approach to explore the evolutionary space available to interacting proteins.

## Results

### Design and generation of nonsynonymous *PKR* and *K3L* genetic variants

Screening pairs of variant alleles first required the generation of single, nonsynonymous variants of both *PKR* and *K3L*. For *PKR*, we designed all variants accessible with a single nucleotide polymorphism (SNP) in four windows of interest (Figure 2A); these windows contain residues in the tertiary structure that interface with eIF2a and residues under positive selection (Figure 2B) (Elde et al., 2009; Jacquet et al., 2022; Rothenburg et al., 2009). These windows of interest are defined as Window 1: G255-F278, Window 2: K371-R385, Window 3: S448-M455, and Window 4: E480-I506. This selection criteria amounted to 374 missense variants, 31 nonsense variants, and an additional 4 synonymous variants (D260, G264, K371, V484) for a total of 409 variants across Windows 1-4 (Figure S1).

**Figure 2.**
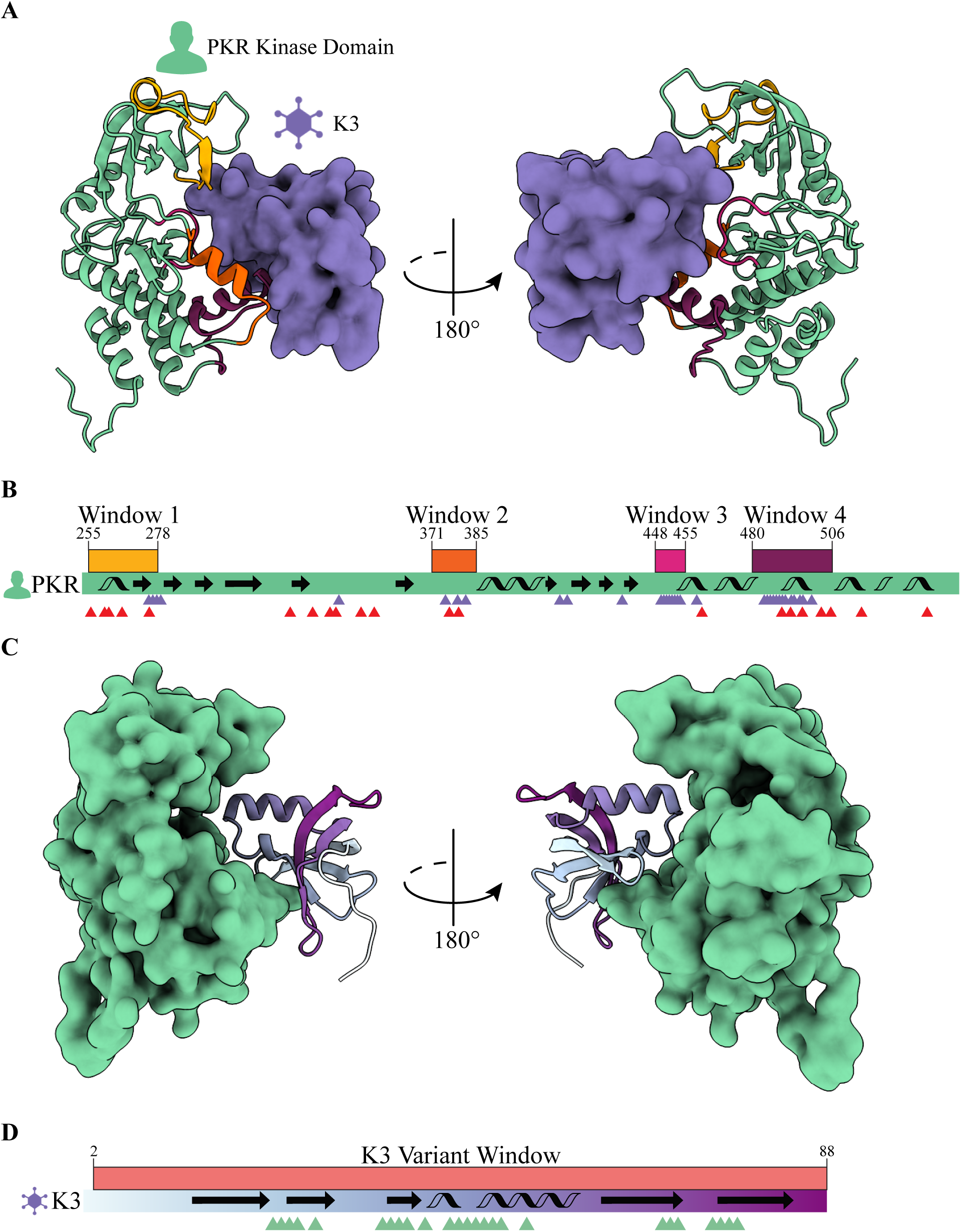
Genetic variants of PKR and K3 were generated in select windows of interest within each protein. **(A)** PKR variants were made across four windows of interest in the PKR kinase domain that interface with the VACV pseudosubstrate antagonist K3. The predicted PKR-K3 complex was generated by aligning PDB 1LUZ Chain A to PDB 2A1A Chain A using PyMol (see Methods). **(B)** *PKR* variants were made across four windows of interest in the PKR kinase domain that interface with K3 and are under positive selection. The PKR kinase domain coding sequence is depicted (green rectangle) with four windows of interest above in yellow, orange, magenta, and burgundy. Secondary structural elements are indicated with arrows (beta sheets) and helices (alpha helices). Purple and red triangles denote PKR residues that contact K3 and are under positive selection, respectively. K3 contact sites are defined as PKR residues within 5 Å of the predicted PKR-K3 complex (see Methods). **(C)** *K3L* variants were made across the entirety of the protein (residues 2-88). (D) *K3L* variants were made across a single window of interest that spanned all residues except the start codon (2-88). The K3 coding sequence is depicted (rectangle with cyan to purple gradient) with a single variant window above in red. Secondary structural elements are indicated with arrows (beta sheets) and helices (alpha helices). Green triangles denote K3 residues that contact PKR as defined by a 5 Å distance (see Methods).

In parallel, we designed a library of variants to explore the local evolutionary space available for K3 to antagonize PKR (Figure 2C). Because K3 is a smaller protein than PKR we were able to design all SNP accessible nonsynonymous variants across the entirety of *K3L* (from VACV strain Western Reserve) between start and stop codons (L2-Q88, Figure 2D). This amounted to 461 missense variants, 35 nonsense variants, and an additional 4 synonymous variants (C5, I39, H47, M84) for a total of 500 variants (Figure S2).

To make the *PKR* and *K3L* variant libraries we designed variant primer tile sets. When used to PCR-amplify the target gene, each variant primer introduces a select, single-residue variant into the resulting gene fragment (PCR-1, Figure S3). Multiple variant primers were pooled together in a single PCR reaction, such that the final PCR product contained a mix of gene fragments, each with a single variant. The reverse barcode primer of the PCR reaction was at the end of the target gene and contained a stretch of randomly mixed nucleotides (Figure S4) such that each variant gene fragment was followed by a unique nucleotide sequence (Figure S5A). To build whole-length genes from the gene fragment (PCR-1), each variant tile set PCR reaction was paired with a separate PCR reaction (PCR-2) that contained the rest of the target gene, the cloning vector sequence, and homology arms that complemented the PCR-1 fragment. These two fragments were then combined via Gibson Assembly to generate a pool of select variants with unique barcodes. 15 variant primer tile sets were designed for the four windows of interest in the PKR kinase domain (Figure S5B) and 18 variant primer tile sets were designed across the entirety of K3 (Figure S5C).

### Long read sequencing of *PKR* and *K3L* variant libraries

Long read sequencing of the *PKR* and *K3L* variant libraries was done to link each nonsynonymous variant to its unique nucleotide barcode. All designed variants were identified for *PKR* and *K3L* except for PKR-H483Q. An average of 43 barcodes were linked to each *PKR* variant, with an average read depth of 22 reads per barcode (Figure S6, Figure S7). 90% of the PacBio reads mapped to intended, single-residue, nonsynonymous variants with another 2% mapping to WT and synonymous variants (Figure S8). For *K3L* a mean of 62 barcodes were linked to each designed variant, with an average read depth of 18 reads per barcode (Figure S9, Figure S10). 83% of the PacBio reads mapped to intended, single-residue, nonsynonymous variants with another 1% mapping to WT and synonymous variants (Figure S11).

### Combining the *PKR* and *K3L* variant libraries to form an all-pairs library

*PKR* and *K3L* variant libraries were combined after generating nonsynonymous variant libraries. The combination of 409 *PKR* variants with 500 *K3L* variants creates a total of 204,500 unique pairs. We combined the *PKR* and *K3L* libraries by digesting the plasmids to produce a “donor” *K3L* fragment and a “receiver” *PKR* fragment (Figure S12). Importantly, the fragments contained homology arms such that Gibson Assembly of the *K3L* “donor” fragment into the *PKR* “receiver” plasmid placed the barcodes adjacent to one another, forming a “paired barcode” that could be read via Illumina short-read sequencing. The two fragments contain 20 base pair (bp) homology arms for Gibson Assembly and were transformed into electrocompetent cells, plated on LB-AMP, recovering ∼2.5 million colonies per all-pairs library replicate such that each unique variant pair had approximately 10 barcode replicates.

### Yeast growth assay to screen PKR and *K3L* all-pairs library

We leveraged a yeast growth assay to screen combinations of *PKR* and *K3L* variants (Kawagishi-Kobayashi et al., 1997). In this assay, galactose-induced expression of PKR inhibits protein synthesis and slows yeast growth (Figure S13). However, co-expression of PKR and K3 results in growth recovery due to K3 antagonism of PKR. Attaching unique barcode sequences to each *PKR* and *K3L* variant allowed use to measure PKR and K3 functionality with the yeast growth assay *en masse*. A pool of yeast cells was transformed with the plasmid library encoding combinations of *PKR* and *K3L*. We induced expression of PKR and K3 and isolated plasmid DNA from surviving cells at 0 and 8 hours post induction. The region of the plasmid encoding the paired barcodes was PCR-amplified and sequenced on an Illumina MiSeq short-read sequencer, from which we calculated the abundance of each paired barcode over the time series to characterize the interaction between PKR and K3 variants. Importantly, each *PKR* and *K3L* variant in a cell could be identified from the paired barcode read by referencing the PacBio sequencing results. Using this readout, the paired barcodes representing a functional PKR that discriminates against K3 would decrease in abundance over time, and the paired barcodes representing functional K3 variants with robust antagonism of PKR would increase over time.

### Variants of opportunity and constraint were identified in *PKR*

Results from the yeast growth assay highlight variants of opportunity and constraint in *PKR*. PKR functional scores were calculated from the barcode fold changes between timepoints (see Methods). As expected, all nonsense variants generated across the PKR kinase domain were characterized as having reduced function relative to WT PKR (Figure 3A). Additionally, six of eight *PKR* variants previously characterized as having enhanced discrimination against WT K3 had positive PKR functional scores (Figure 3B, highlighted in blue) (Seo et al., 2008). WT PKR had a functional score of 0.06, with a mean missense functional score of -0.03. A total of 149 *PKR* variants displayed improved evasion of K3 antagonism over WT PKR while maintaining kinase function. Newly identified variants with positive PKR functional scores, such as I270M, E375R, L482F, E490Q, and I506M, were found across all four windows of interest and are highlighted in blue (Figure 3A). Most variants in window 3 had negative PKR functional scores, and window 3 variants overall had significantly lower scores than variants from windows 1, 2, and 4 (Figure 3B). Window 3 is distinct in that (1) it contains part of the kinase active site, (2) is primarily composed of residues that contact both K3 and eIF2α (see Methods), and (3) there are no residues under positive selection.

**Figure 3.**
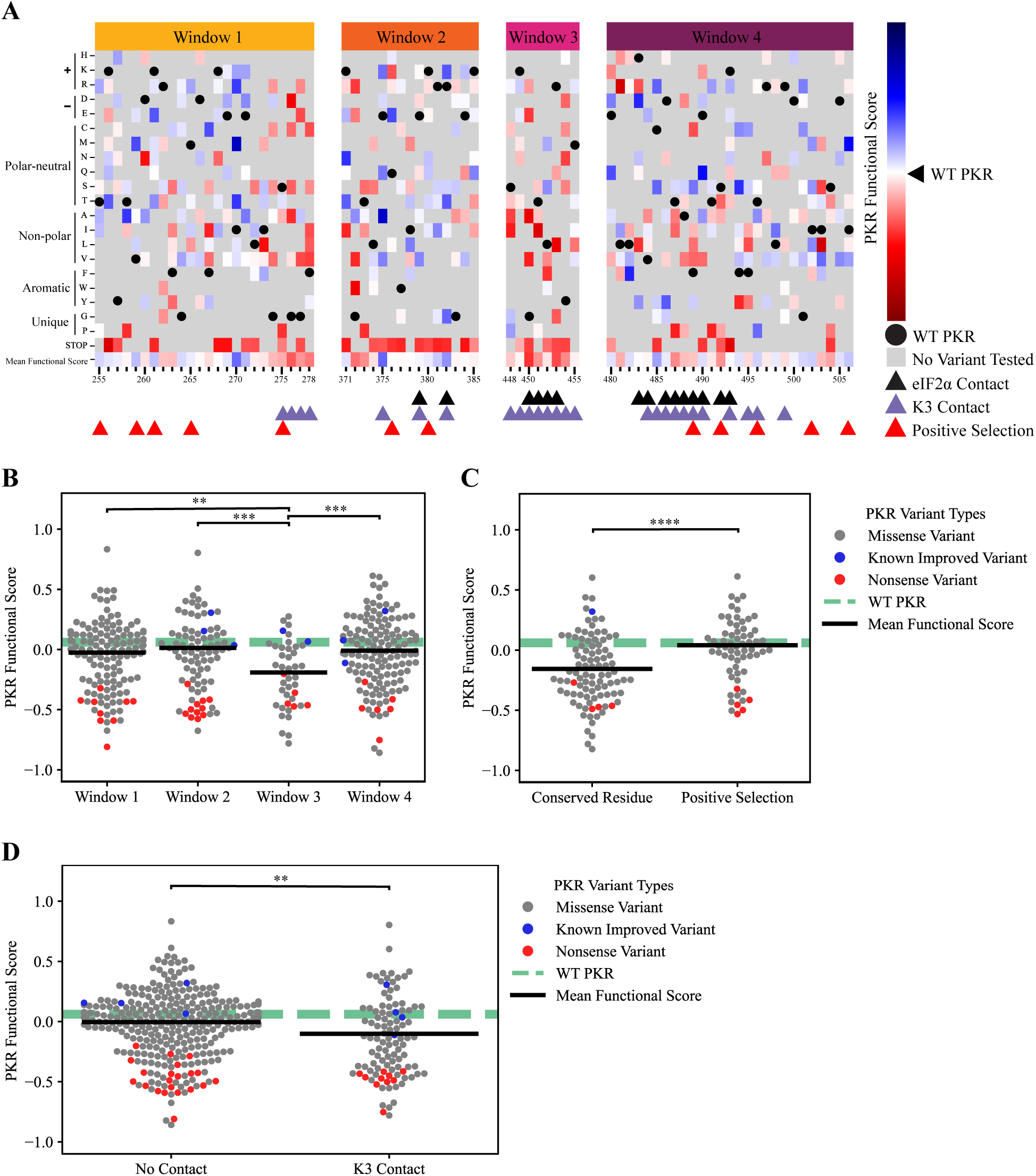
Variants of opportunity and constraint for PKR to discriminate against K3 and phosphorylate eIF2α. **(A)** PKR functional scores (see Methods) of *PKR* variants across the four windows of interest in the kinase domain. Blue cells denote variants with improved function to phosphorylate eIF2α in the presence of WT K3, as seen through the inhibition of translation and yeast growth in our assay (see Figure S13). Red cells denote *PKR* variants with impaired function compared to WT PKR, either unable to evade K3 or are nonfunctional and do not phosphorylate eIF2α. White cells indicate that the variant performs similarly to WT PKR, with gray cells being untested variants. Triangles below the heatmap indicate which PKR residues contact eIF2α (black), contact K3 (purple), or are under positive selection among primates (Elde et al., 2009) (red). Black circles are WT PKR residues, for reference. K3 and eIF2α contact sites are defined as PKR residues within 5 Å of the predicted PKR-K3 or PKR-eIF2α complex (see Methods). Window 1 = G255-F278, Window 2 = K371-R385, Window 3 = S448-M455, Window 4 = E480-I506. **(B)** We found a statistically significant difference in variant functional scores separated into windows of interest. (One-way ANOVA, F(2)=5.4, p=.001). A Tukey post-hoc test revealed significant pairwise differences between Window 3 and Windows 1, 2, and 4 (p<.004, .001, .001). Markers depict functional scores for PKR missense variants (gray), known improved variants (blue), and nonsense variants (red). The WT PKR functional score is shown for reference (dashed green line), and the mean functional score for missense variants in each grouping is shown (black bar). Nonsense mutations were not included when calculating the mean and statistical significance between groups. Window 1 = G255-F278, Window 2 = K371-R385, Window 3 = S448-M455, Window 4 = E480-I506. **(C)** The difference in mean PKR functional scores between *PKR* variants that do not contact K3 (M=-.005, SD=0.255) and sites that contact K3 (M=-.102, SD=0.318) is significant (Independent T-test, t(372)=2.99, p=0.003). Markers depict functional scores for PKR missense variants (gray), known improved variants (blue), and nonsense variants (red). The WT PKR functional score is shown for reference (dashed green line), and the mean functional score for missense variants in each grouping is shown (black bar). K3 contact sites are defined as residues within 5 Å of the predicted PKR-K3 complex (see Methods). Nonsense mutations were not included when calculating the mean and statistical significance between groups. **(D)** The difference in mean PKR functional scores between *PKR* variants at conserved residues (M=-.15, SD=0.28) and residues under positive selection (M=0.04, SD=0.23) is significant (Independent T-test, t(148)=-4.46, p=1e-05). Markers depict functional scores for PKR missense variants (gray), known improved variants (blue), and nonsense variants (red). The WT PKR functional score is shown for reference (dashed green line), and the mean functional score for missense variants in each grouping is shown (black bar). Conserved residues are maintained among all four eIF2α kinases (Dar et al., 2005) and residues under positive selection are defined by Elde et al. (2009). Nonsense mutations were not included when calculating the mean and statistical significance between groups. **p<0.01, ***p<0.001, ****p<.0001

We compared functional scores between *PKR* variants at conserved and rapidly evolving sites under positive selection. We found a significant difference between variants located at conserved residues between the four ISR kinases and PKR residues under positive selection across 21 primate species (Figure 3C) (Elde et al., 2009). We also compared the functional scores of *PKR* variants located at contact sites and found a significant difference between PKR variants at sites that do and do not contact K3 (Figure 3D). Contact sites were defined as PKR residues within 5 Å of the K3 in the predicted complex (see Methods), with contact sites identified across all four windows of interest.

Overall, 149 *PKR* variants had functional scores greater than WT PKR (Figure 3A). These variants were distributed across all 4 windows of interest but were predominant in Windows 1, 2, and 4 (Figure 3B), with enrichment at PKR residues under positive selection (Figure 3C). Residues I270, E375, R382, and C485 harbor multiple variants that evade K3 antagonism. These residues were not found to be under positive selection when comparing PKR homologs across primate species (Elde et al., 2009) but I270, E375, and R382 were found to be under positive selection in vertebrate species (Rothenburg et al., 2009). 256 *PKR* variants had lower functional scores than WT PKR (Figure 3A) and were found among all four windows of interest with enrichment at the conserved Window 3 containing the PKR kinase active site. Lower functional scores were also moderately enriched at contact sites with K3 (Figure 3D).

### Variants of opportunity and constraint were identified in *K3L*

K3 functional scores (see Methods) from the yeast growth assay also highlight variants of opportunity and constraint across the entirety of K3 (Figure 4A), with a WT K3 functional score of -.06 and a mean missense variant functional score of -0.18 (Figure 4B). 104 variants were identified with higher K3 functional scores over WT K3, including newly identified variants L30V, E37A, A41E, E53K, and M84V. The majority the nonsense mutations had reduced function in comparison to WT K3, with three exceptions at the C-terminus and another three located at K16*, E37*, and K64*. Consistent with past reports, we identified K3-H47R as an improved variant and K3-H47Q, H47D, and K74A having reduced functional scores in comparison to WT K3 (Dar & Sicheri, 2002; Kawagishi-Kobayashi et al., 1997; Seo et al., 2008). In comparing functional scores of variants at sites that contact PKR no significant difference was observed (Figure 4C). However, we did observe that the few *K3L* variants that made K3 more like eIF2α resulted in a mildly significant increase in the K3 functional score (Figure 4D).

**Figure 4.**
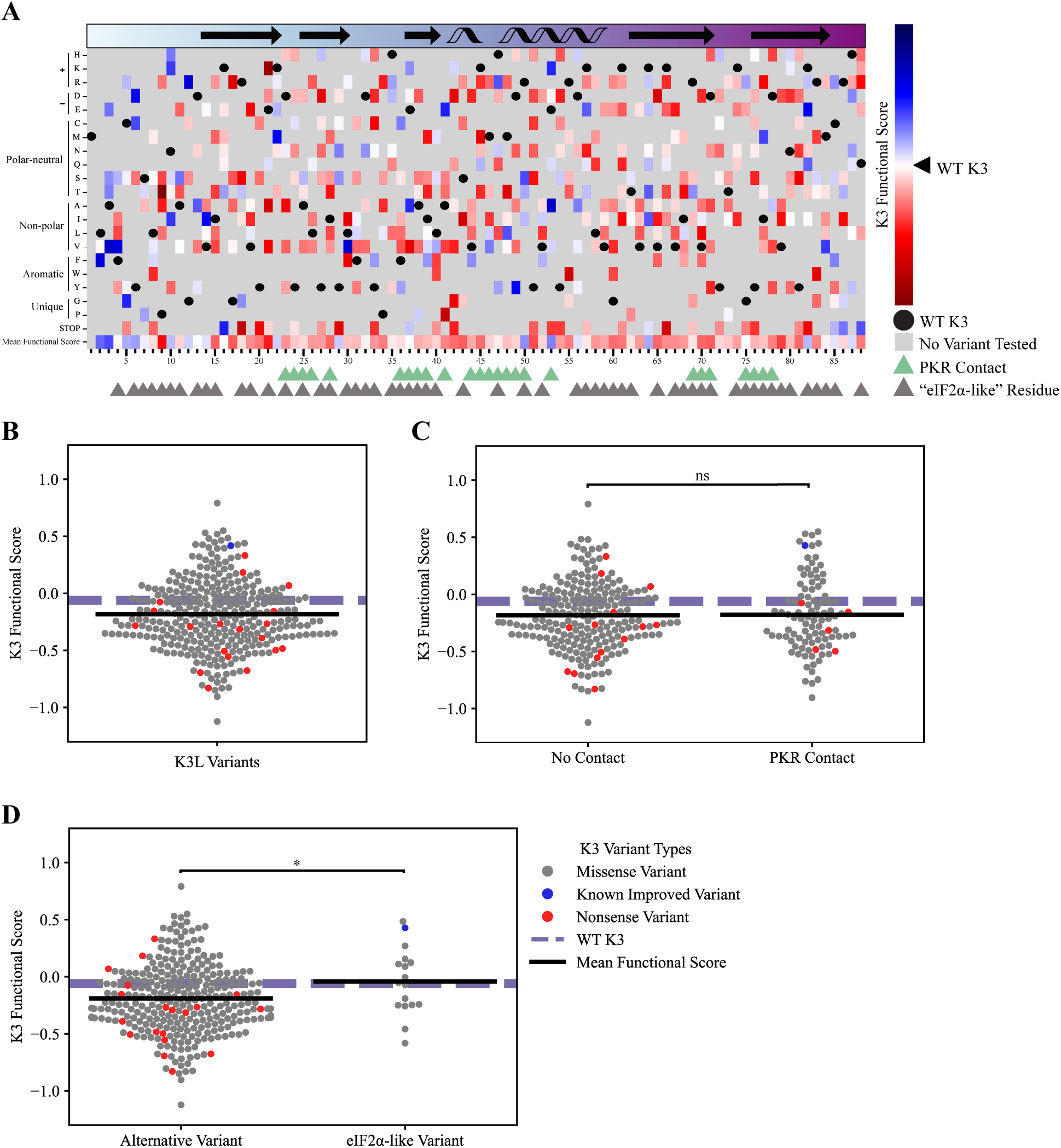
Variants of opportunity and constraint for K3 to antagonize PKR. **(A)** K3 functional scores (see Methods) for variants across the entirety of K3 (L2-Q88). Blue cells denote variants with improved function to antagonize WT PKR, red cells denote *K3L* variants with impaired function compared to WT K3. White cells indicate that the variant performs similarly to WT K3, with gray cells being untested variants. The secondary structure of K3 is displayed in the rectangle above the heatmap with arrows (beta sheets) and helices (alpha helices). The coloring of the rectangle matches the color gradient used for the K3 structure in Figure 2C-D. Triangles below the heatmap indicate contact residues with PKR (green) and grayscale triangles indicate residues that align to eIF2α (see Methods). Black circles are WT K3 residues, for reference. **(B)** The full range of K3 functional scores for *K3L* highlight variants of opportunity and constraint. Markers depict functional scores for *K3L* missense variants (gray), known and nonsense variants (red). A single known improved variant, K3 H47R, is depicted in blue. The WT K3 functional score is shown for reference (dashed purple line), and the mean functional score for missense variants is shown (black bar). Nonsense mutations were not included when calculating the mean functional score. **(C)** No significant difference in K3 functional scores between variants that do and do not contact PKR. The difference in mean K3 functional scores between *K3L* variants the do and do not contact PKR was not found to be significant. Markers depict functional scores for *K3L* missense variants (gray), known and nonsense variants (red). The WT K3 functional score is shown for reference (dashed purple line), and the mean functional score for missense variants in each grouping is shown (black bar). PKR contact sites are defined as residues within 5 Å of the predicted PKR-K3 complex (see Methods). Nonsense mutations were not included when calculating the mean functional score. **(D)** A mildly significant difference in the mean K3 functional scores was observed between *K3L* variants that make K3 more similar to the protein sequence of eIF2α (M=0.99, SD=0.20) when compared to all other *K3L* variants (M=0.90, SD=0.19, Independent T-test, t(299)=-1.97, p=0.05). Markers depict functional scores for *K3L* missense variants (gray), known and nonsense variants (red). The WT K3 functional score is shown for reference (dashed purple line), and the mean functional score for missense variants in each grouping is shown (black bar). eIF2α-like variants are defined based on protein sequence alignment between K3 and eIF2α (see Methods). Nonsense mutations were not included when calculating the mean functional score. *p<.05, ns=not significant

Overall, 104 *K3L* variants had functional scores greater than WT K3 and were distributed across the entirety of the coding sequence, including the alpha-helix insert (H47-V59) that contains K3-H47R. 216 *K3L* variants had reduced functional scores in comparison to WT K3, including a majority of the nonsense variants as well as K3-H47Q, H47D, and K74A (Kawagishi-Kobayashi et al., 1997; Seo et al., 2008).

## Discussion

### Functional variants identified at sites under positive selection

Our results both recapitulate the findings of previously characterized *PKR* variants and identify many novel variants. 149 *PKR* variants displayed improved evasion of K3 antagonism over WT PKR while maintaining kinase function, including the previously characterized variants M455V, E480D, E375V, I378T, T491S, and S448G (Seo et al., 2008) and the newly identified variants I270M, E375R, L482F, E490Q, and I506M. For *K3L*, 104 variants were identified with higher K3 functional scores over WT K3, including the previously characterized H47R variant, and were found across the entirety of the protein sequence.

Our results underscore the role that PKR residues under positive selection play in evading K3 antagonism (Elde et al., 2009; Rothenburg et al., 2009). We found that PKR residues under positive selection were enriched for not only functional but improved *PKR* variants that evade K3 and maintain functionality. In contrast, residues conserved among the four ISR kinases did not tolerate variation. For instance, missense variants across the kinase domain active site (Window 3) were found to disrupt protein function.

These findings emphasize the functional plasticity of PKR at residues under positive selection and expand the identification of functional variants. We found many beneficial variants in Windows 1, 2, and 4, including PKR’s alpha helix D and alpha helix G, which were previously identified as regions under positive selection that alter PKR’s sensitivity to K3 (Elde et al., 2009; Rothenburg et al., 2009; Seo et al., 2008). However, we additionally found many variants in the N-lobe of the kinase domain that also allow PKR to evade K3 yet are not nearly as close to the interaction site, specifically in Window 1. These functional variants are in residues under positive selection or in proximity to them. It is possible that these variants alter the dimerization and activation of PKR (Dar et al., 2005; Dey et al., 2005)and must be characterized in the absence of K3.

### Defining an “improved” K3 antagonist

We identified 104 *K3L* variants that display improved antagonism of PKR over WT K3, including the previously characterized variant H47R (Kawagishi-Kobayashi et al., 1997) as well as newly identified variants L30V, E37A, A41E, E53K, and M84V. Defining an “improved” K3 is difficult. It is possible that a *K3L* variant with milder inhibition of PKR could increase the dissemination of the virus and be considered and “improved” variant. For example, the myxoma virus elicits very different sequela in brush rabbits and European rabbits. Brush rabbits are the natural hosts of myxoma virus, and the virus causes relatively benign, localized lesions (Kerr & Best, 1998; Regnery & Miller, 1972). The myxoma virus protein M156 is a highly effective inhibitor of brush rabbit PKR, preventing both a shutdown in translation as well as stimulating an immune response (Yu et al., 2022). In contrast, myxoma virus protein M156 is a moderate inhibitor of European Rabbit PKR (Yu et al., 2022). This moderate inhibition stunts cellular translation but also stimulates pro-inflammatory cytokines, making myxoma virus highly virulent in European rabbits (Kerr & Best, 1998; Regnery & Miller, 1972). This response may promote the systemic spread of the virus, as leukocytes have been shown to play an essential role in the viruses dissemination (Fenner & Woodroofe, 1953; Kerr, 2012).

Given this example, an improved K3 would moderately inhibit PKR to increase dissemination of the virus. However, emerging myxoma isolates carry a M156 L98P variant which abolishes inhibition of European rabbit PKR (Peng et al., 2016). Given this evidence it is difficult to define an improved form of M156. Our results find that there is plenty of room for K3 to improve or reduce its antagonism of WT PKR, which may allude to a tunable form.

### Methodological strategies to explore evolutionary space

Our experimental approach is noteworthy for two methodological achievements. First, we designed and generated single residue, nonsynonymous variants of both *PKR* and *K3L* in a pooled format. We used a PCR-based strategy featuring variant primers to generate hundreds of variants of each gene in a pooled format. We aimed to link each variant to approximately 10 unique DNA barcodes that were identified by long-read sequencing. This approach allows us and others to systematically generate hundreds of specific genetic variants for unbiased screens, and that variant information can be compressed and localized to the barcode sequence using long-read sequencing. This compression step enables both high-throughput screening and the opportunity to combine variant alleles together, as we did with *PKR* and *K3L*. Finally, the existence of multiple barcodes per variant resulted in a pool with internal biological replicates.

Second, we merged two variant libraries into an extensive all-pairs library in which every *PKR* variant was paired with every *K3L* variant. This all-pairs library contained over 200,000 unique variant pairs, each with approximately 10 replicate barcode sequences. Our cloning strategy to combine variants resulted in a paired barcode in which the *PKR* and *K3L* barcodes are adjacent. We could then use short-read sequencing to track variant pairs over time, allowing the yeast growth assay to accommodate hundreds of thousands of variant combinations in a single flask. We expect that continued advances in sequencing technology will permit ever-larger combinatorial libraries to be screened, allowing broader evolutionary spaces to be explored.

This experimental approach can be pushed even further to characterize all-pairs of *PKR* and *K3L*. We expect this approach can be broadly applied towards other protein-protein interactions of interest, including other evolutionary arms races.

### Evolutionary insights and future directions

Genetic variation is a continuous source for innovation that is filtered by selection, and in the case of positive selection these variations are often retained. The ability to proactively explore and characterize the local variant space is a step towards anticipating future variation. This advance offers an opportunity to incorporate resilient features into a myriad of designs from agricultural crops to antiviral therapeutics. In the context of an evolutionary arms race, in which two proteins are rapidly adopting variation, we can further extend this approach to evaluate the robustness of each variant against all other variants, which is a future direction of this project enabled by the *PKR*-*K3L* all-pairs library.

This work emphasizes positive selection as a strong signal for pliable sites within proteins, though not all variants at these sites are functional. The evolutionary arms race between PKR and K3 continues to offer insights into the resilience of the immune system. These findings extend our understanding of protein evolution and offer new directions for future research.

## Methods

### Design of YCp50-RD plasmid

We designed a multiple-cloning site (MCS) into the single copy plasmid YCp50 to generate the plasmid YCp50-RD (receiver-donor), which was used for subsequent cloning and combining *PKR* and *K3L* variants. The MCS contained seven unique restriction enzyme cut sites (AgeI, BsiWI, BstEII, NotI, MluI, PvuII, and SacI), was designed such that the eventual combination of *PKR* and *K3L* variants would be combined using restriction enzyme cut sites with 20 bp homology arms for Gibson Assembly (Gibson et al., 2009), with *PKR* and *K3L* barcodes being separated by a 20 bp spacer. YCp50 was digested using restriction enzymes EcoRI-HF (NEB Cat#R3101S) and BspDI (NEB Cat#R0557S) with rCutSmart buffer, followed by purification and size selection on 1% agarose gel with .6 µg/mL ethidium bromide (BioRad Cat#1610433), selecting for the 7,961 bp band. The extracted band was purified using a QIAquick Gel Extraction (Qiagen Cat#28706) and eluted with 30 µL of water. For Gibson Assembly the purified band was combined with a single-stranded DNA oligonucleotide containing the MSC and homology arms spanning the digest site, named Oligo 1 (Table S1), in a 1:5 molar ratio in a total volume of 5 µL, along with an additional 5 µL 2x Gibson Assembly Master Mix (NEB Cat#E2611L). The reaction mixture was incubated at 50°C for 1 hour. 1 µL of the Gibson Assembly reaction was transformed into 10 µL 5-alpha competent E. coli (NEB Cat#C2987I) and plated onto 10 cm LB-AMP plates (IPM Scientific Cat#11006-016) and incubated overnight at 37°C. Colonies were selected for overnight outgrowth in 3 mL LB-AMP (IPM Scientific Cat#11006-004), followed by a QIAprep spin miniprep (Qiagen Cat#27106) and Sanger sequencing to validate the plasmid sequence using the primer Oligo 2. This produced the plasmid YCp50-RD.

### Cloning *PKR* and *K3L* into YCp50-RD

We cloned WT PKR and WT K3 into the landing pad of YCp50-RD to produce the YCp50-WT_PKR and YCp50-WT_K3 plasmids, respectively. The plasmid p1419 (Kawagishi-Kobayashi et al., 1997) contains *PKR* under the control of the galactose-inducible pGAL/CYC1 promoter to control expression of PKR in yeast; we PCR-amplified the expression construct using primers Oligo 3 and Oligo 4 to place the *PKR* expression cassette between landing pad cut sites AgeI and BstEII. Oligo 4 incorporated a synthetic terminator for *PKR* (Tsynth1) (Curran et al., 2015), a 28 bp nucleotide spacer sequence, and a 26 bp barcode; both primers added a 20 bp homology arms to the YCp50-RD plasmid. pGAL/CYC1-PKR was amplified using the polymerase PfuUltraII (Agilent Technologies Cat#600674) for 30 cycles with an annealing temperature of 55°C, followed by gel purification. YCp50-RD was digested with restriction enzymes AgeI-HF (NEB Cat#R3552S) and BstEII-HF (NEB Cat#R3162S), followed by gel purification. The digested YCp50-RD and pGAL/CYC1-PKR fragments were joined via Gibson Assembly in a 1:5 molar ratio, followed by transformation into 5-alpha competent *E. coli* and plating onto LB-AMP. A resultant clone, validated by Sanger sequencing, was designated YCp50-WT_PKR.

To produce YCp50-WT_K3, the *K3L* coding sequence was amplified from the plasmid pC140 (Kawagishi-Kobayashi et al., 1997) using primers Oligo 5 and Oligo 6. The TDH3 promoter, also known as pTDH3 or pGPD, was amplified from genomic DNA of the yeast strain BY4742 (Brachmann et al., 1998) using primers Oligo 7 and Oligo 8. Oligo 5 incorporated a synthetic terminator for K3 (Tsynth8) (Curran et al., 2015), a 28 bp nucleotide spacer sequence, a 26 bp barcode, and a 20 bp homology arm to the YCp50-RD. Oligo 6 incorporated a correction to the *K3L* sequence from pC140 to match the VACV WT strain *K3L* sequence with a Val2Leu substitution, and provided a 20 bp homology arm to the end of pTDH3. WT K3 was paired with pTDH3 for constitutive expression in yeast. pTDH3 and *K3L* were amplified using PfuUltraII, followed by gel purification. YCp50-RD was digested with restriction enzymes NotI-HF (NEB Cat#R3189S) and PvuII-HF (NEB Cat#R3151S), followed by gel purification and extraction. The digested YCp50-RD pTDH3, and K3 fragments were joined via Gibson Assembly in a 1:5:5 molar ratio, followed by transformation into 5-alpha competent *E. coli* and plating onto LB-AMP. A resultant clone, validated by Sanger sequencing, was designated YCp50-WT_K3.

### Design of nonsynonymous variant primers for *PKR* and *K3L*

Doped primers were designed and grouped into variant primer tile sets to generate select nonsynonymous variants across windows of interest in the *PKR* and *K3L* coding sequence. Doped primers, or mixed bases, allow for select bases to be randomly incorporated into oligos at specific positions. Select bases are designated using IUPAC degenerate base symbols symbols (e.g. “D” is an equal mix of “A”, “G”, and “T”). We used custom Python (v3.9.6) scripts to design each primer with a 20 bp homology arm, variant region with a doped codon to generate nonsynonymous variants flanked by doped codons generating synonymous variants using IUPAC degenerate base symbols, and a priming sequence with a melting temperature of 50°C (Cornish-Bowden, 1985; Guido & Drake Jr, 2009). To explore local evolutionary space, we selected nonsynonymous variants accessible with a single nucleotide polymorphism (SNP) using *PKR* (Ensemble ENSG00000055332, CDS EIF2AK2-001) and *K3L* (Vaccinia virus Western Reserve strain, Genbank AY243312.1, location 27305:27572(-)) coding sequences. Primers were designed to generate nonsynonymous variants across the four windows of interest in PKR (G255-F278, K371-R385, S448-M455, and E480-I506) and from residues L2-Q88 in K3. Primers were synthesized by Integrated DNA Technologies, then were pooled into variant primer tiles sets, with 15 sets for *PKR* (Oligos 10-24) and 18 sets for *K3L* (Oligos 25-42). The variant primer tile sets were paired with barcode primers, one each for *PKR* and *K3L* (Oligo 43 and Oligo 44, respectively). Each barcode primer consisted of a 20 bp homology arm matching the sequence downstream of the gene, a barcode region with twenty fully doped bases, and a priming sequence with a melting temperature of 55°C. Mixed bases were limited to 5 nucleotide stretches to prevent the unintended generation of restriction sites, spacing these stretches with “AA” or “TT” sequences (Liu et al., 2019).

We also designed primer pairs to amplify the remainder of the target gene along with all plasmid components. These involved a constant forward primer for *PKR* and *K3L*, each consisting of a 20 bp homology arm to the respective barcode primer and a priming sequence with a melting temperature of 55°C. A reverse primer was designed for each variant primer tile set to provide homology to the homology arm of the forward primers, with 15 reverse primers for *PKR* and 18 reverse primers for *K3L*.

### Generation of *PKR* and *K3L* variant libraries

We made *PKR* and *K3L* variant libraries by Gibson Assembly of two PCR amplicons generated using the primers described above. Each Gibson Assembly reaction involved two PCR amplicons, referred to as PCR-1 and PCR-2. Each PCR-1 was generated using a variant primer tile set and a barcode primer. For the *PKR* variant library, 15 PCR-1 reactions were completed using the variant primer tile sets (Oligos 10-24) paired with the *PKR* barcode primer Oligo 43 to amplify from YCp50-WT_PKR plasmid using PfuUltraII polymerase with an annealing temperature of 45°C. Each reaction generated a pool of *PKR* gene fragments containing select nonsynonymous variants paired with a unique barcode sequence. In parallel, 15 PCR-2 reactions were completed using Oligo 45 paired with Oligos 47-61 to amplify from YCp50-WT_PKR plasmid using Herculase II polymerase (Agilent Technologies Cat#600677) using cycling conditions for vector targets >10 kilo-base pairs, each reaction generated a larger vector fragment with 20 bp homology arms complementing one of the PCR-1 amplicons. PCR-1 and PCR-2 amplicons were digested with DpnI (NEB Cat#R0176S) for 2 hours at 37°C to remove the YCp50-WT_PKR template, followed by gel purification. Each PCR-1 amplicon was paired with its corresponding PCR-2 amplicon for Gibson Assembly in a 1:5 molar ratio for a total of 15 reactions.

For the *K3L* variant library, 18 PCR-1 reactions were completed using the variant primer tile sets (Oligos 25-42) paired with the *K3L* barcode primer Oligo 44 to amplify from the YCp50-WT_K3 plasmid using PfuUltraII polymerase with an annealing temperature of 45°C. Each reaction generated a pool of *K3L* gene fragments containing select nonsynonymous variants paired with a unique barcode sequence. 18 PCR-2 reactions were completed using Oligo 46 paired with Oligos 62-79 to amplify from the YCp50-WT_K3 plasmid using Herculase II polymerase. Each reaction generated a larger vector fragment with 20 bp homology arms complementing one of the PCR-1 amplicons. PCR-1 and PCR-2 amplicons were digested with DpnI to remove the YCp50-WT_K3 template, followed by gel purification. Each PCR-1 amplicon was paired with its corresponding PCR-2 amplicon for Gibson Assembly in a 1:5 molar ratio for a total of 18 reactions.

2 µL of each Gibson Assembly were transformed into 50 µL of 5-alpha competent *E. coli* and all cells were plated on LB-AMP and incubated overnight at 37°C. Colonies were counted on each plate and plates were bottlenecked at 30x colonies per selected nonsynonymous variant (i.e. each nonsynonymous variant would be linked to approximately 30 unique barcode sequences). Plates were washed with 15 mL LB-AMP, cell densities were measured at an OD of 600, and 6 ODs from each reaction were pooled to form a single *PKR* variant library and a single *K3L* variant library. 1 mL glycerol stocks were made for each library and a 200 mL LB-AMP outgrowth was grown to a concentration of 3 OD600/mL followed by a Qiagen MAXI plasmid prep (Qiagen Cat#12963).

### PacBio sequencing of barcoded variant libraries

Both the *PKR* and *K3L* variant libraries were sequenced using the PacBio Sequel II instrument to identify the barcodes that were linked to each nonsynonymous variant. 16 µg of each variant library was digested with two restriction enzymes to create equally sized fragments of approximately 2100 bp for sequencing, AgeI-HF/NotI-HF were used for the *PKR* variant library and PvuI-HF/BamHI-HF for the *K3L* variant library. Digests were incubated for 2 hours at 37°C then purified using a QiaQuick PCR purification kit (Qiagen Cat#28106). Fragments of 2100 bp were size-selected using a SageELF instrument (Sage Science) before PacBio Circular Consensus Sequence (CCS) HiFi sequencing on a Sequel II instrument (Pacific Biosciences). From the CCS reads we generated a table of *PKR* and *K3L* barcodes paired with genetic variants using alignparse v0.2.6 (Crawford & Bloom, 2019).

### Generation of all-pairs library of PKR and K3 combinations

We merged the PKR and K3 variant libraries such that yeast transformed with the single copy plasmid would express a single variant of PKR and a single variant of K3. Each variant library was digested with two restriction enzymes to generate linear DNA fragments with 20b bp complementary homology arms for Gibson Assembly. 10 µg of the *PKR* variant library was digested with 5 µL of NotI-HF and 5 µL PvuII-HF in a 50 µL reaction to generate a linear vector fragment of approximately 10,270 bp. 10 µg of the *K3L* variant library was digested with 5 µL of BstEII-HF and 5 µL SacI-HF (NEB Cat#R3156S) in a 50 µL reaction to generate a linear insert fragment of approximately 1130bp. Both fragments were gel purified. The *PKR* and *K3L* fragments were combined in three separate Gibson Assembly reactions to generate three replicate libraries using a 1:5 molar ratio with 100 ng of the *PKR* vector in a reaction volume of 20 µL and incubated at 50°C for 1 hour. A standard ethanol precipitation was performed on each reaction, adding 100 µL of 100% ETOH and incubating overnight at -20°C before resuspending the pellets in 2 µL of water (Sambrook & Russell, 2006). Note that the use of restriction enzymes to produce the components of the Gibson Assembly reaction was preferred over PCR to avoid template switching, which could scramble the linkage between barcodes and nonsynonymous variants.

We transformed each of the three concentrated reactions into *E. cloni* 10G SUPREME electrocompetent cells (Lucigen Cat#60080-2) using a MicroPulser Electroporator (BioRad), with 2 µL of the concentrated reaction combined with 25 µL of competent cells and splitting the electroporation onto two 15 cm LB-AMP plates. 1 µL of competent cells was plated with 100 µL of water on a 10 cm plate to estimate the total colony count on the two 15 cm plates, which produced an estimate of approximately 2 million colonies per electroporation. Plates were washed with 25 mL LB-AMP, combining the two plates for each of the three reactions to form three replicates of the all-pairs library. 1 mL glycerol stocks were made and each replicate library was expanded in 200 mL LB-AMP to a concentration of 3 OD600/mL, followed by a Qiagen MAXI plasmid prep.

### Screening the all-pairs library in a yeast growth assay

We transformed each of the three *PKR*-*K3L* all-pairs libraries into yeast and induced PKR expression, sampling the three replicate cultures over time. The all-pairs plasmid libraries were transformed into the yeast strain BY4742 (Brachmann et al., 1998) using a standard large-scale high efficiency lithium acetate transformation protocol (Gietz & Schiestl, 2007) and plated onto 15 cm CSM-Ura (Sunrise Science Products Cat#1004-010) 2% dextrose plates, using 1/1,000 and 1/10,000 dilutions on 10 cm plates to estimate colony counts. Plates were washed with 25 mL CSM-Ura 2% dextrose, combined into pools, and 1 mL glycerol stocks were made for each of the replicates.

To start the yeast growth assay, replicates were seeded at .001 OD600/mL in 330 mL CSM-Ura 2% dextrose and grown overnight for 16 hours at 30°C, 200 revolutions per minute (rpm). The following morning a first timepoint sample (designated as “0 hours”) was taken. 15 ODs of cells were harvested from each replicate flask and spun down in a 1.5mL Eppendorf tube at 5,000 rpm for 1 minute, after which supernatant was removed and the pellet was frozen at - 80°C. 40 ODs of cells from the remaining cultures were pelleted in 50 mL conical tubes at 3,000 rpm, supernatant was removed, and cells were resuspended in 200 mL CSM-Ura 2% galactose (.2OD600/mL) to induce PKR expression. A second timepoint was taken at 8 hours by pelleting 15 ODs at 3,000 rpm in a 50 mL conical tube, then resuspending cells in 1 mL CSM-Ura 2% galactose and pelleting in a 1.5 mL Eppendorf tube at 5,000 rpm for 1 minute, removing supernatant, and freezing the pellet at -80°C. With two timepoints (0 hours and 8 hours) and three replicates, a total of six samples were taken through the yeast growth assay.

### Plasmid extraction and barcode amplification

To quantify fluctuations of barcode abundance between timepoints in the yeast growth assay we harvested plasmids from the two timepoints, then amplified and sequenced the adjacent *PKR* and *K3L* barcodes in each plasmid. Plasmids were harvested from the yeast timepoint samples using a modified QIAprep spin miniprep protocol (Singh & Anthony Weil, 2002). Sampled cell pellets of 15 OD were first thawed at room temperature, followed by the addition of 250 µL QIAprep P1 buffer and 2 µL zymolase (2.5 units/µL), and incubation at 37°C for 30 minutes, followed by adding 250 µL P2 buffer and following the manufacturer instructions for the remainder of the protocol.

We paired *PKR* and *K3L* barcodes by amplifying from each of the six samples using a two-step PCR protocol, the first to amplify the paired barcode region from the plasmids and the second to attach Illumina adapters and Nextera indices for downstream sequencing and demultiplexing sample reads. For PCR reaction 1, five forward and five reverse primers contain Nextera transposase adapter sequences (Illumina Document#1000000002694) “TCGTCGGCAGCGTCAGATGTGTATAAGAGACAG” (read 1, forward primers) or “GTCTCGTGGGCTCGGAGATGTGTATAAGAGACAG” (read 2, reverse primers) for tagmentation, a staggered region of 0-4 doped “N” bases for adding sequence diversity, and a priming sequence with a melting temperature of 60°C, forming Oligos 80-89 (Wu et al., 2015). The five forward and five reverse primers were pooled together to form Oligo 90 and Oligo 91. 1 µL of each primer Oligo 90 and Oligo 91 were combined with 50 ng of sample plasmid DNA and 25 µL Kapa Hifi Hotstart ReadyMix (Kapa Biosystems Cat# KK2602), and topped off with water to a total volume of 50 µL. The PCR reaction 1 thermocycler is as follows: (1) 95°C for 3 minutes, (2) 98°C for 20 seconds, (3) 65°C for 15 seconds, (4) 72°C for 1 minute, repeat steps 2-4 for a total of 18 cycles, (5) 72°C for 1 minute, (6) 12°C hold. PCR products were purified using a MinElute PCR purification kit (Qiagen Cat#28006) using 11 µL of water for the final elution, then quantified using a Qubit dsDNA High Sensitivity (HS) kit (ThermoFisher Scientific Cat#Q32851).

For PCR reaction 2, 10 ng of amplicon DNA from PCR reaction 1 was combined with 10 µL of Nextera adapter index primers (Illumina Cat#20027213) and 25 µL Kapa HiFi Hotstart ReadyMix, and topped off with water to a total volume of 50 µL. The PCR reaction 2 thermocycler protocol is as follows: (1) 100°C for 45 seconds, (2) 100°C for 15 seconds, (3) 60°C for 30 seconds, (4) 72°C for 30 seconds, repeat steps 2-4 for a total of 8 cycles, (5) 72°C for 1 minute, (5) 12°C hold. PCR products were purified using a MinElute PCR purification kit (Qiagen Cat#28006) using 11 µL of water for the final elution, then quantified using a Qubit dsDNA Broad Range (BR) kit (ThermoFisher Scientific Cat#Q32850). 500 ng of the PCR reaction 2 amplicons were gel purified on a Size Select II E-Gel (ThermoFisher Scientific Cat#G661012) for approximately 13 minutes to extract the 260bp amplicon band of interest, followed by quantification with a Qubit dsDNA HS kit. Amplicon samples were diluted to 4 nM before being pooled together, followed by manufacture denature and dilution protocols (Illumina Document#15039740v10) before sequencing on an Illumina MiSeq instrument.

### PKR and K3 functional scores and screening analysis

We extracted *PKR* and *K3L* barcode sequences from the Illumina reads and mapped the barcodes back to their corresponding select nonsynonymous variants using the table of *PKR* and *K3L* barcodes paired with genetic variants generated from the PacBio CCS HiFi reads. Paired reads were assembled into contiguous sequences using PEAR v0.9.11 (Zhang et al., 2014), followed by Bartender v1.1 (Zhao et al., 2018) to extract and cluster *PKR* and *K3L* barcode sequences using the respective barcode search patterns “CAAGG[25-27]GGTGA” and “GCCGC[25-27]GACAG”.

We wrote Python v3.9.6 scripts to tally paired barcodes and map them back to *PKR* and *K3L* genetic variants using the table generated from the PacBio CCS HiFi reads. Barcode counts were grouped by genetic variant and normalized to the total paired barcode count for each of the six samples, followed by a fold change of the normalized count from timepoint 1 (0-hour) to timepoint 2 (8-hour) for each of the three replicates, then taking the mean fold change across the three replicates for each variant. Variants were filtered for all *PKR* variants paired with WT K3 and all *K3L* variants paired with WT PKR, excluding variant-variant pairs. Variants with a mean total read count of <18 reads across the three replicates at timepoint 1 (0 hours) were dropped from further analysis, which filtered down to 387 *PKR* variants and 333 *K3L* variants.

For *PKR* variants paired with WT K3 we assigned a functional score (FSPKR) defined as:

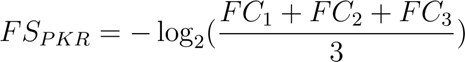

where FC is the fold change of a *PKR* variant paired with WT K3 in each of the three replicates. Of note, the log2 of the mean fold change is inverted such that *PKR* variants that decreased in abundance were assigned positive functional scores, as increased PKR activity in evading K3 antagonism and phosphorylating eIF2α would be expected to be associated with decreasing relative abundance in the yeast growth assay, resulting in a fold change less than 1. For *K3L* variants paired with WT PKR we assigned a functional score (FSK3) defined as:

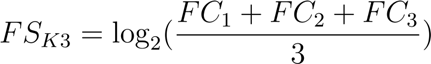

where FC is the fold change of a *K3L* variant paired with WT PKR in each of the three replicates. Here, the log2 of the mean fold change is not inverted, as *K3L* variants with improved PKR antagonism would be expected to be associated with increasing relative abundance in the yeast growth assay, resulting in a fold change greater than 1.

To define PKR residues contacting K3, the K3 structure PDB 1LUZ was aligned to eIF2α (chain A) of the PKR-eIF2α structure complex (PDB 2A1A) using PyMOL v2.5.4. The following PKR residues within 5 Å of the aligned K3 structure were selected: 276, 277, 278, 335, 337, 375, 382, 450, 451, 453, 454, 485, 486, 487, 488, 489, 490, 491, 492, 493, 496, and 499. Selected residues contained branching side chains within 5 Å of PKR branching side chains. All other PKR residues were categorized as not contacting K3. We also categorized PKR residues that are conserved or are under positive selection. The following PKR residues were categorized as conserved based on their conservation among the four human ISR kinases: PKR, PERK, HRI, and GCN2 (residues 262, 263, 266, 267, 273, 274, 276, 278, 450, 451, 454, 480, 481, 487, 490, 495, 498, and 499) (Dar et al., 2005). The following PKR residues were categorized as being under positive selection when looking across 21 primate PKR sequences: 6, 7, 24, 44, 49, 86, 122, 123, 125, 139, 185, 206, 224, 242, 255, 259, 261, 265, 275, 322, 330, 336, 338, 344, 351, 376, 380, 462, 489, 492, 496, 502, 506, 516, and 538 (Elde et al., 2009).

K3 residues at sites that contact PKR were defined using the same structural alignment used above to define PKR residues that contact K3, selecting for residues within 5 Å of PKR: 22, 23, 24, 25, 27, 36, 37, 38, 39, 41, 44, 45, 46, 47, 48, 49, 50, 53, 69, 70, 71, 75, 76, 77, and 78. All other *K3L* variants were categorized as not having contact with PKR. *K3L* variants that make the protein sequence more like eIF2α were defined based on a MUSCLE v5.1 protein sequence alignment of K3 and eIF2α (PDB 2A1A, Chain A). A nonsynonymous *K3L* variant generating a residue matching the aligned eIF2α residue was defined as eIF2α-like, with all other *K3L* variants being categorized as alternative variants.

## Data access

Supplemental data and code are available under the DOI: 10.5281/zenodo.10183372

## Acknowledgements

We thank Dr. Thomas Dever at the Eunice Kennedy Shriver National Institute of Child Health and Human Development for helpful discussions, experimental guidance for the yeast growth assay, and providing plasmids p1419 and pC140. We thank Dr. Mark Rose of Georgetown University for the plasmid YCp50. We are also grateful to Simone Giovanetti for her helpful feedback and expertise in the lab. This work utilized the computational resources provided by the National Institutes of Health (NIH) HPC Biowulf Cluster (http://hpc.nih.gov) and was supported by the Intramural Research Program of the National Human Genome Research Institute, National Institutes of Health (1ZIAHG200401).

## Supplementary figures

**Figure S1.**
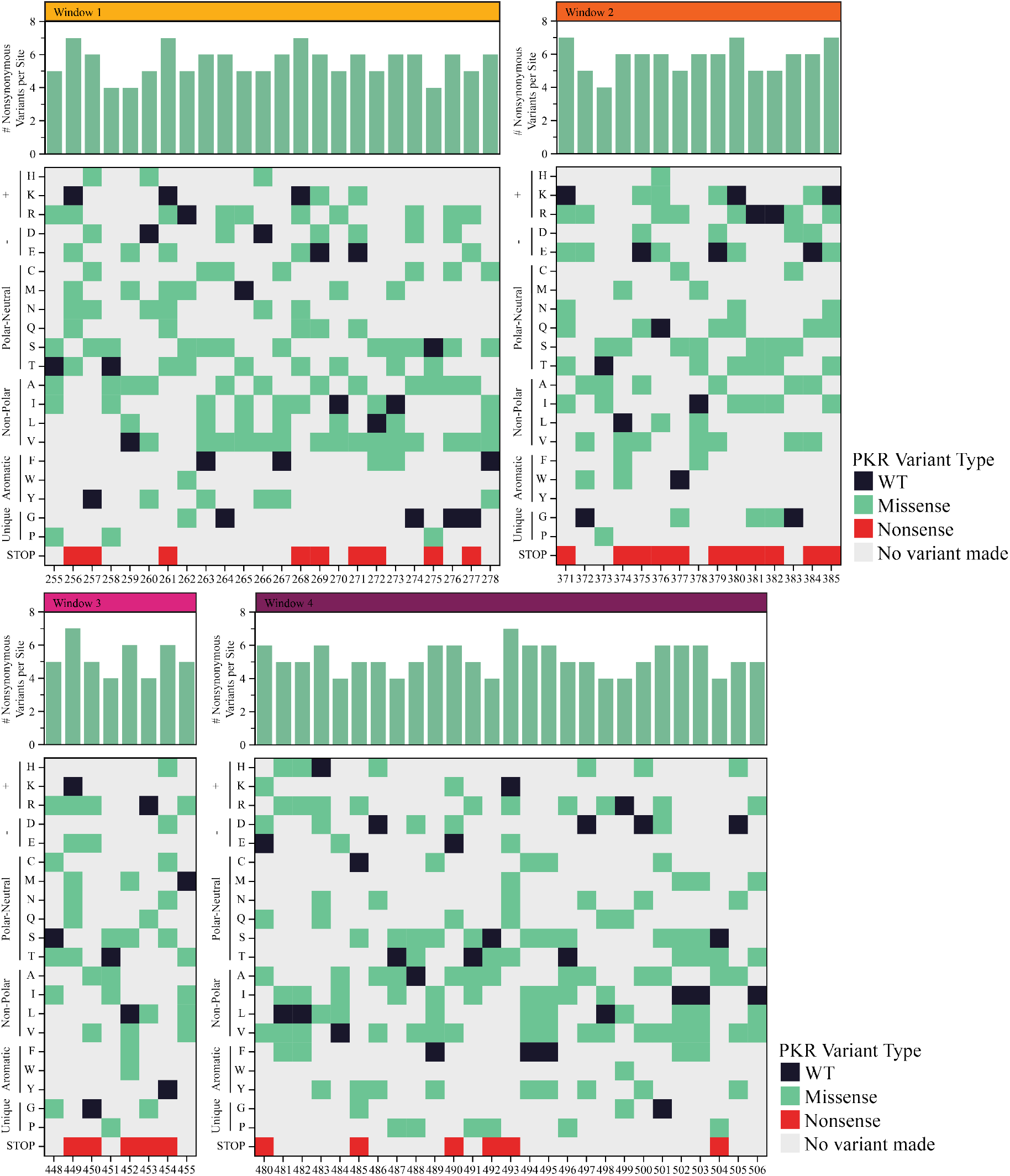
409 nonsynonymous, SNP-accessible variants of interest were selected across four windows of interest in PKR. The top bar chart denotes the total number of variants that were made at each position, ranging from 4-8 variants. The sequence diagram highlights nonsynonymous, SNP-accessible *PKR* variants that were made across the four windows of interest in the PKR kinase domain, with missense variants in green, nonsense variants in red, and variants not made in gray. WT PKR residues are shown in black for reference.

**Figure S2.**
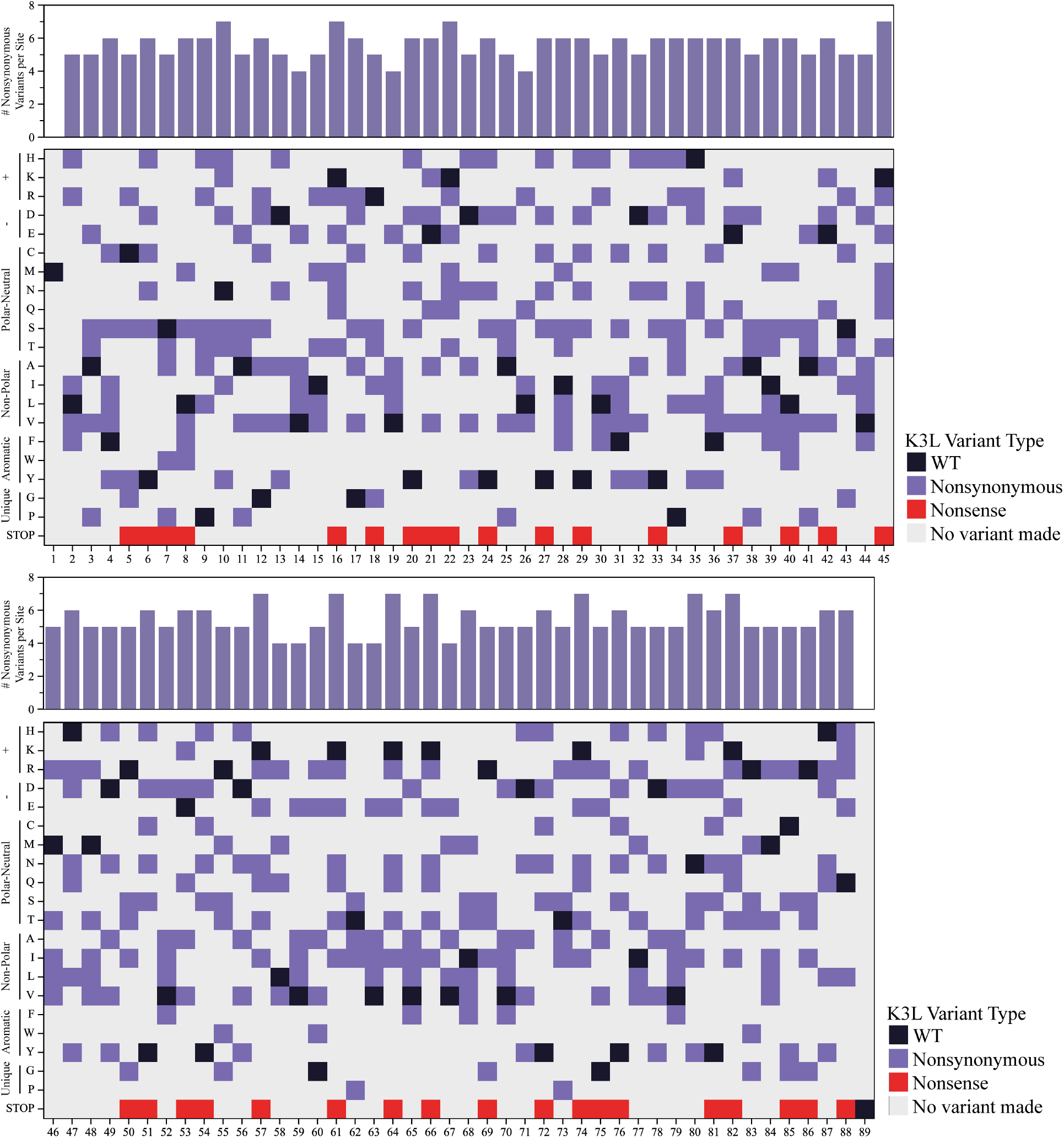
500 nonsynonymous, SNP-accessible variants of interest were selected across the entirety of *K3L*. The top bar chart denotes the total number of variants that were made at each position, ranging from 5-8 variants. The sequence diagram highlights nonsynonymous, SNP-accessible *K3L* variants that were made across the entirety of K3 (L2-Q88), with missense variants in purple, nonsense variants in red, and variants not made in gray. WT K3 residues are shown in black for reference.

**Figure S3.**
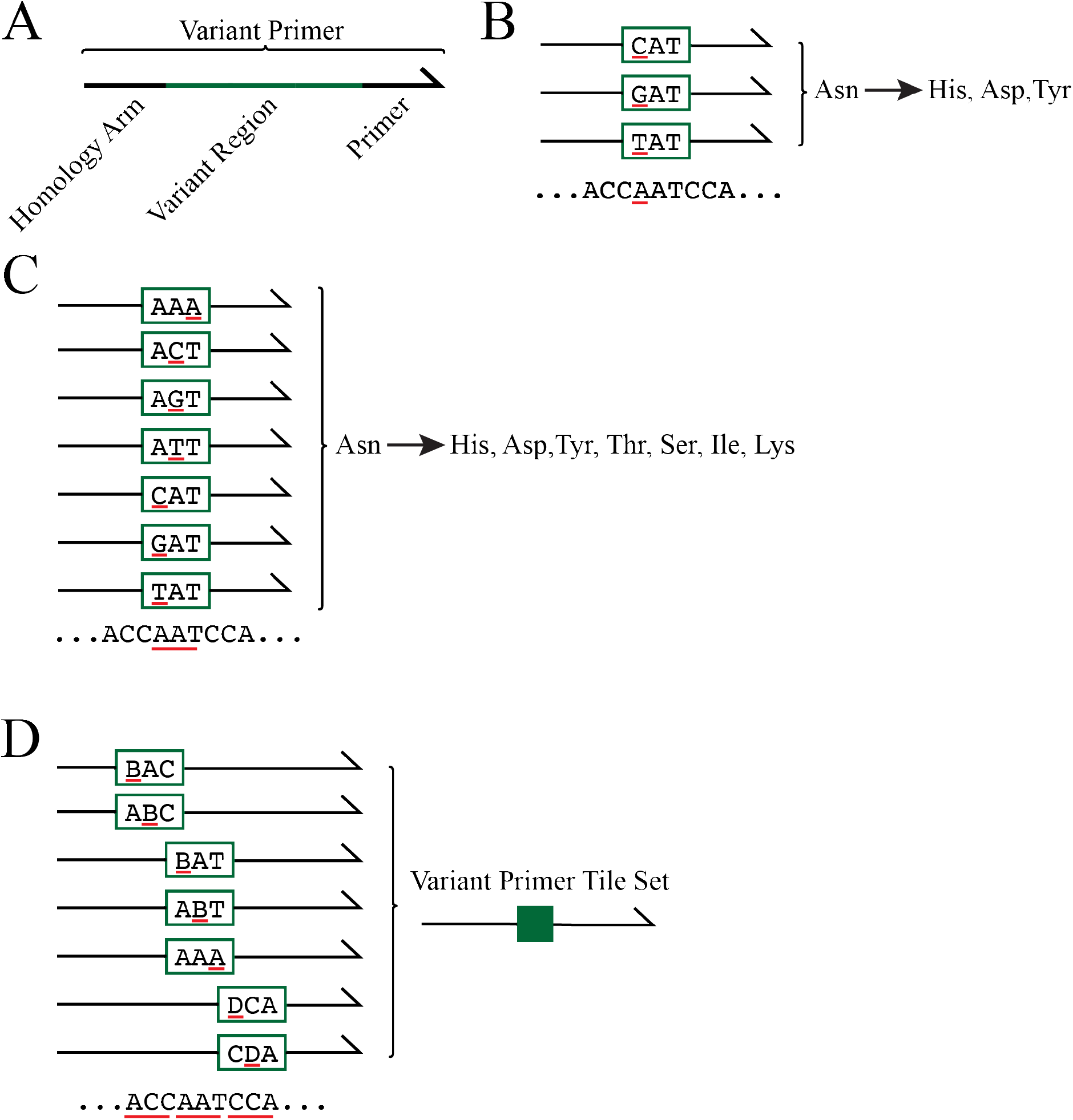
Variant primer tile sets were designed to systematically make single residue nonsynonymous variants. **(A)** Each variant primer is composed of a homology arm, variant region, and primer. **(B)** Nonsynonymous variants are generated by altering the codon in variant region of the primer. This example depicts the codon “AAT” encoding Asn. The first nucleotide in the codon, “A”, is underlined in red, with three codons above having changes to “C”, “G”, and “T” underlined in red, which generate the nonsynonymous variants His, Asp, and Tyr. **(C)** Variant primers were designed across all three nucleotides in each codon, as underlined in red. **(D)** Variant primer tile sets, represented in dark green, were made by pooling variant primers that modify adjacent codons. Primers included in variant primer tile sets have differing variant regions but share homology arms and priming region sequences.

**Figure S4.**
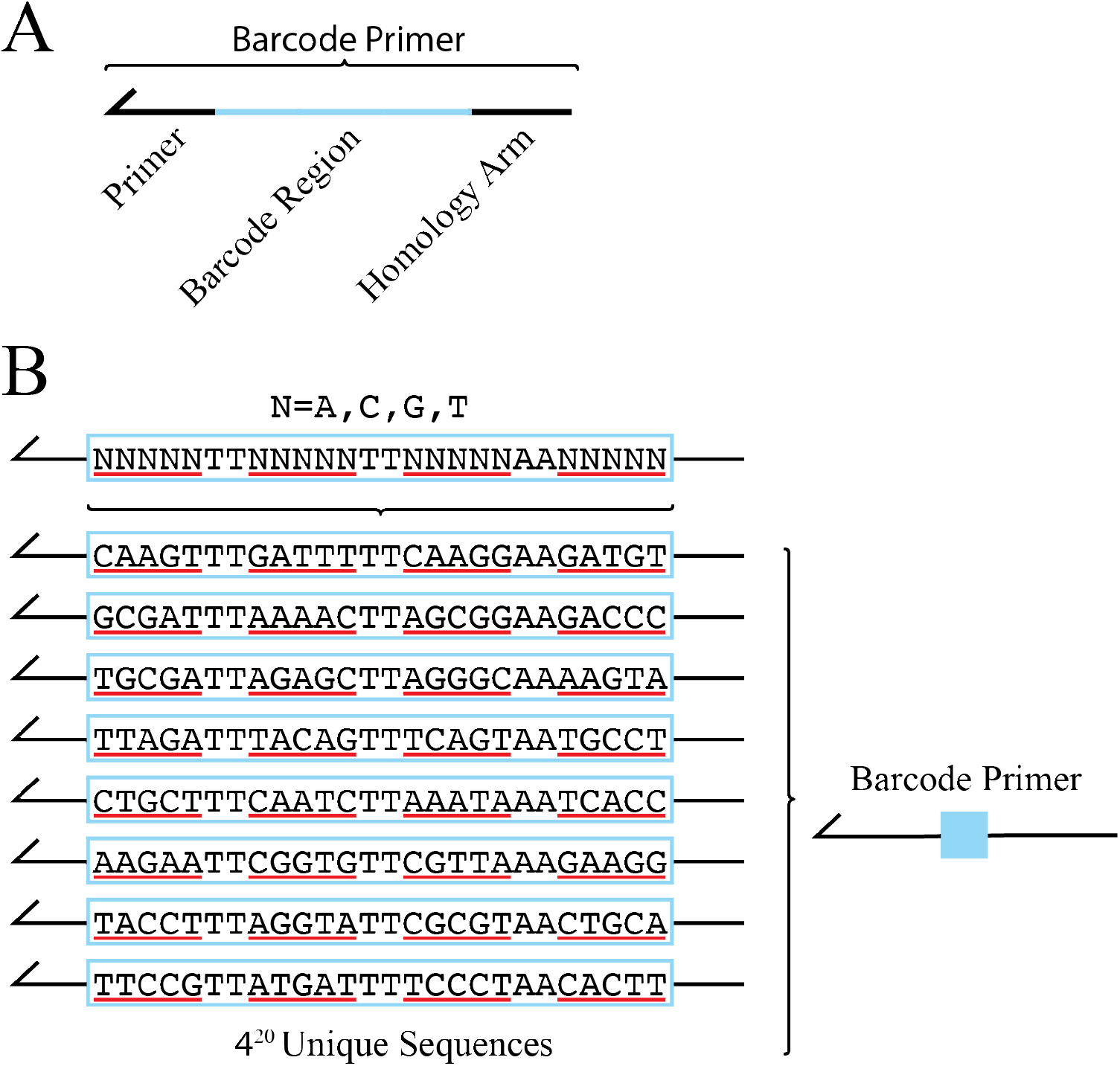
Barcode primers were designed to attach unique nucleotide sequences to each nonsynonymous variant. **(A)** The barcode primer is composed of a homology arm, barcode region, and primer. The barcode primer is paired with each of the variant primer tile sets to attach a unique nucleotide sequence to each nonsynonymous variant made. **(B)** The barcode region of the primer is composed of 20 “N”, each “N” encodes an equal representation of the nucleotides “A”, “C”, “G”, and “T”. A barcode primer with 20 random nucleotides contains 420 unique nucleotide sequences. Dinucleotide sequences “TT” and “AA” are interspersed throughout the barcode region to avoid making unintended restriction enzyme cut sites.

**Figure S5.**
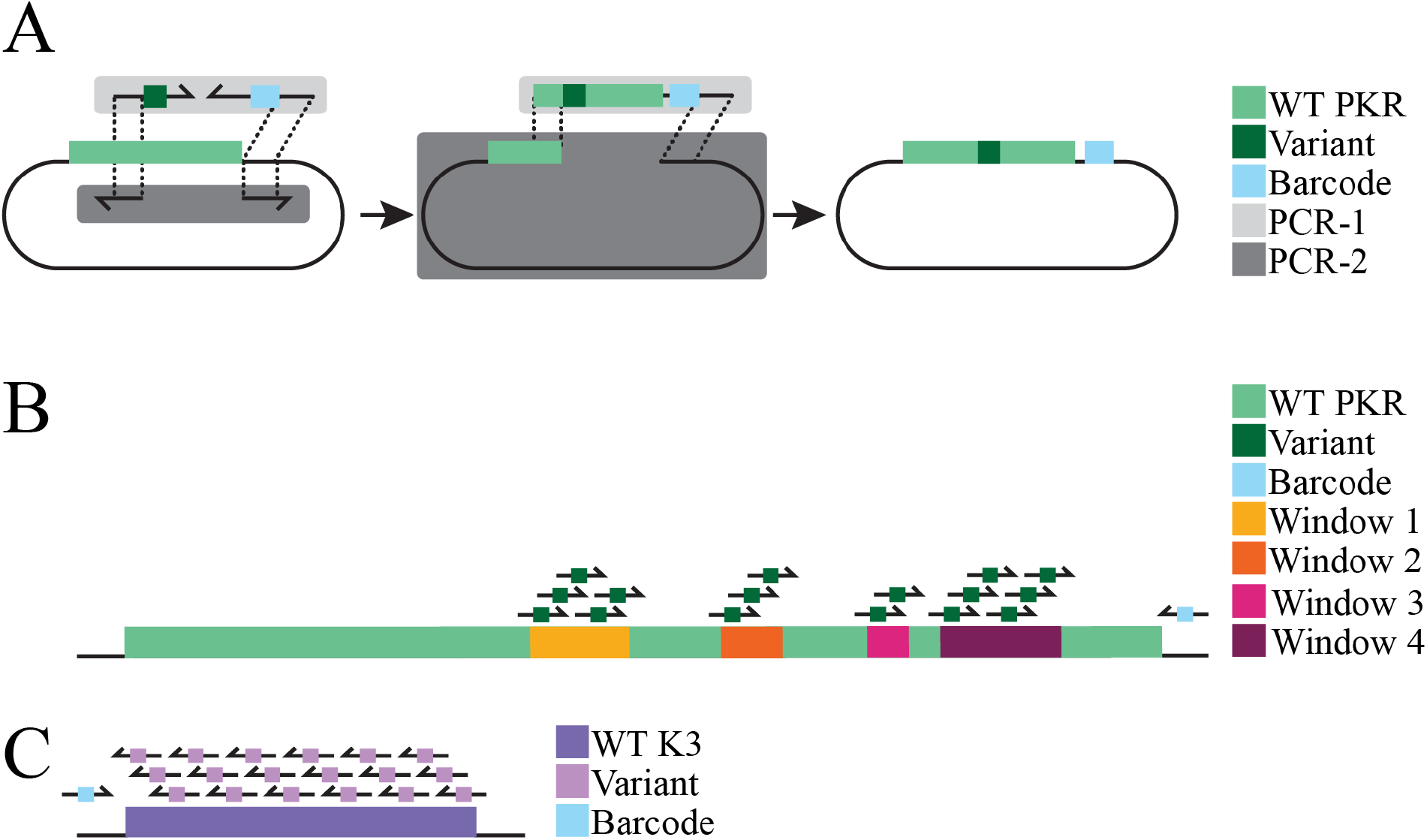
Single residue nonsynonymous variants were PCR amplified and assembled using doped primer tile sets across *PKR* and *K3L*. **(A)** Two separate PCR reactions were used to generate complementary insert and vector fragments. PCR-1 primers (light gray box) include a single variant primer tile set (dark green, see Figure S3D) and a single doped barcode primer (light blue, see Figure S4B) that amplified from WT *PKR* (green) and made the PCR-1 insert fragment containing a select nonsynonymous variant (dark green) and a unique barcode (blue). PCR-2 primers included a single forward and reverse primer that amplified from WT *PKR* and made larger vector fragment with 20 bp homology arms that complement the homology arms of the PCR-1 insert fragment. The two fragments were combined via Gibson Assembly to form a pool of complete vectors, each vector containing a single, nonsynonymous variant with a unique barcode. **(B)** 15 variant primer tile sets were designed to generate select variants across four windows of interest in the *PKR*. The full-length *PKR* sequence is denoted in green, with Windows 1-4 overlaid in yellow, orange, magenta, and burgundy. Variant primer tile sets encode select variants (dark green) and are paired with a single doped barcode primer (blue) for a total of 15 PCR-1 reactions for PKR that generate 435 variants. **(C)** 18 variant primer tile sets were designed to generate select variants across the entirety of *K3L*. The full length *K3L* sequence is denoted in purple; variant primer tile sets encode select variants (pink) and are paired with a single doped barcode primer (blue), making a total of 18 PCR-1 reactions for K3 that generate 527 variants.

**Figure S6.**
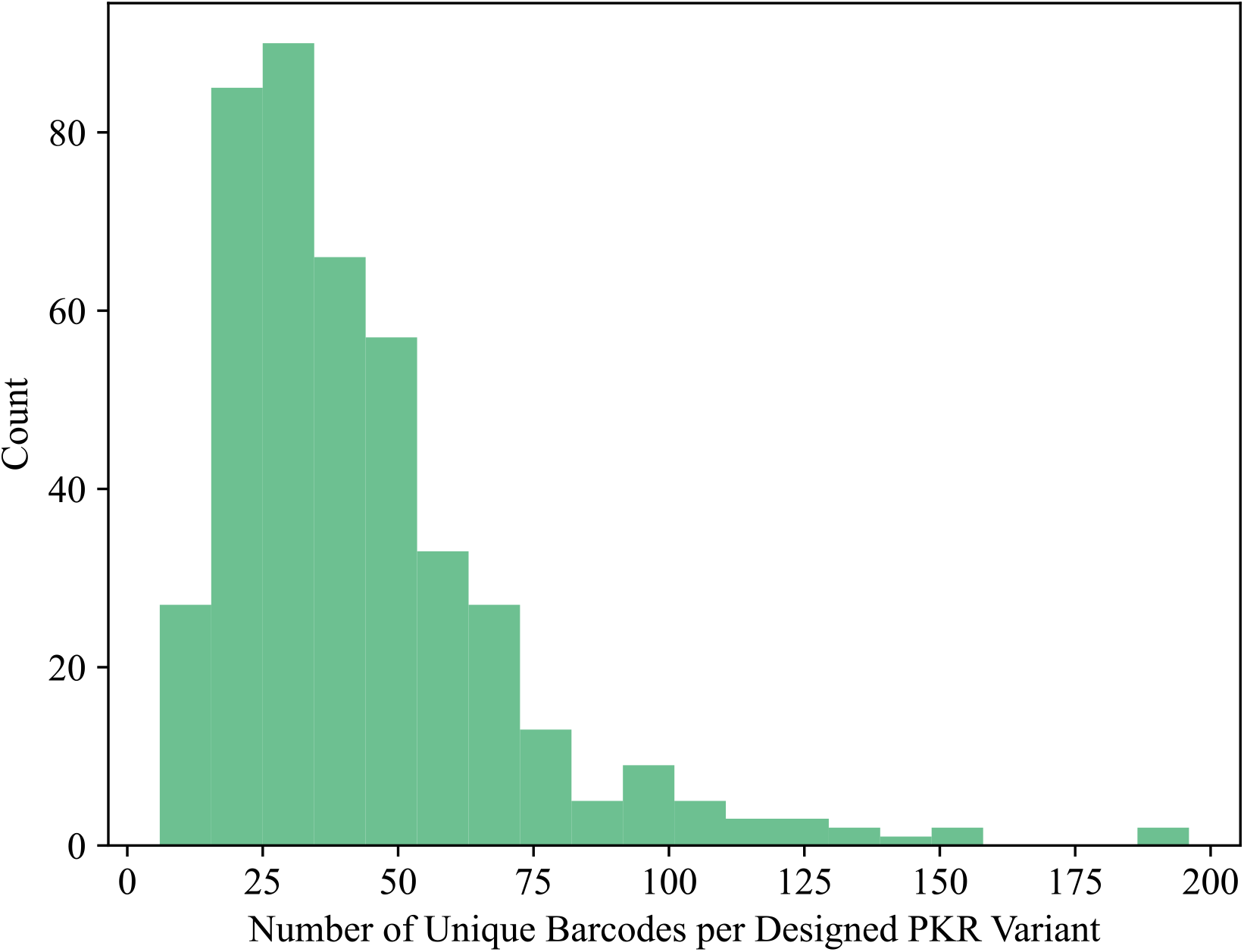
Count of unique barcodes per designed *PKR* variant. Histogram depicts the number of unique barcodes per designed *PKR* variant, with a mean of 43 barcodes per *PKR* variant.

**Figure S7.**
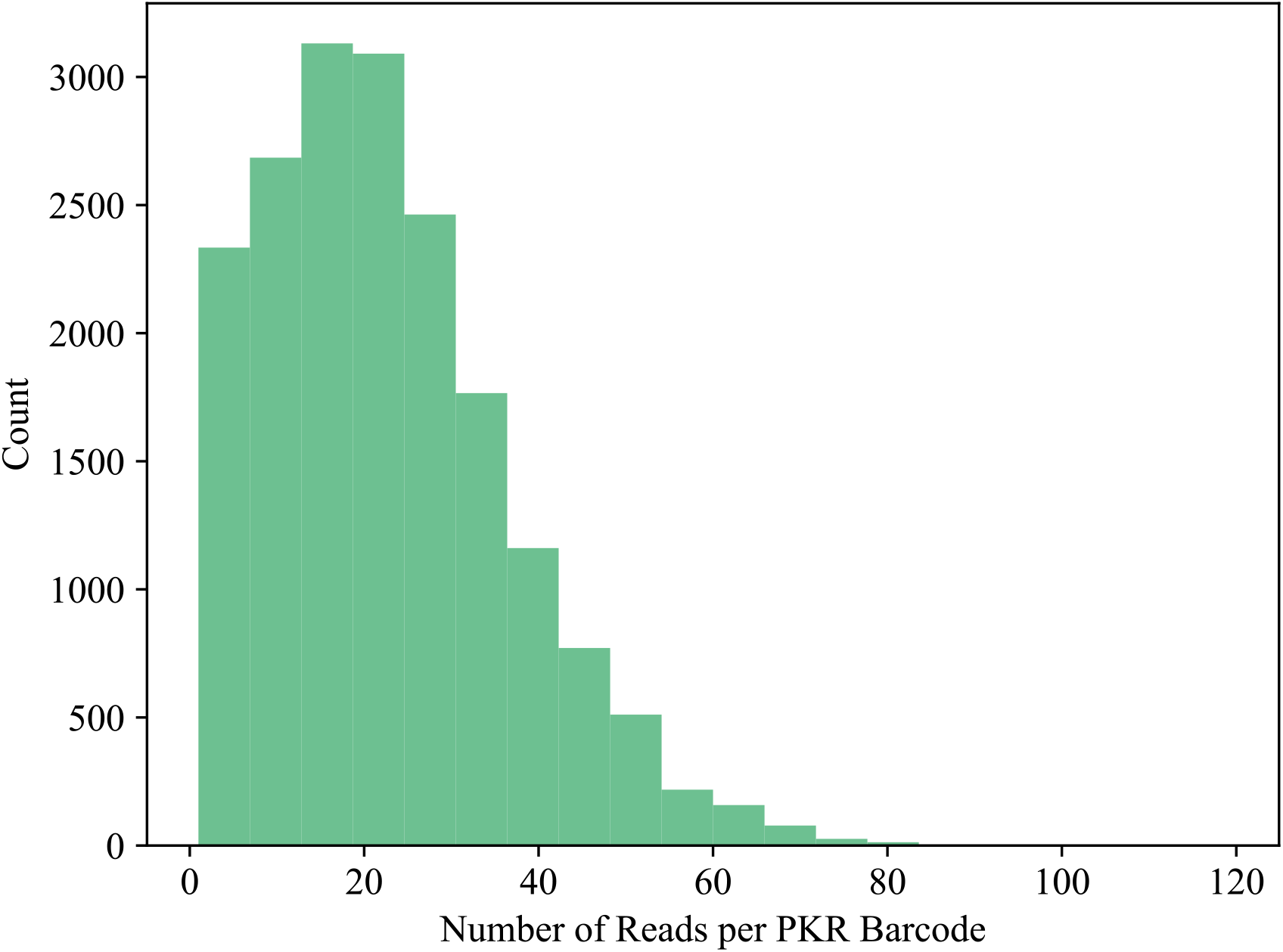
Count of unique barcodes per designed *PKR* variant. Histogram depicts the number of unique barcodes per designed *PKR* variant, with a mean of 43 barcodes per *PKR* variant.

**Figure S8.**
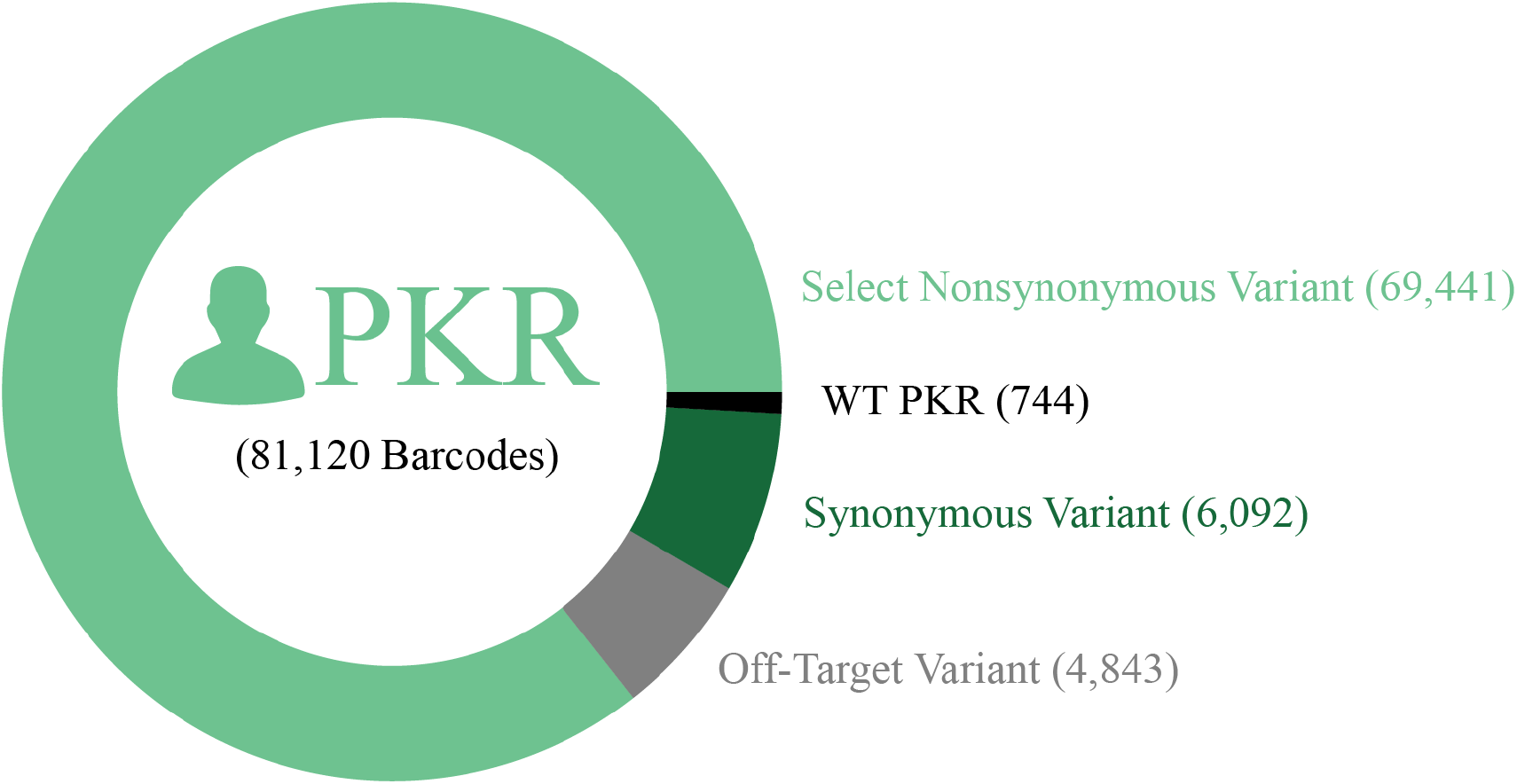
PacBio barcode distribution linked to *PKR* variant types. A total of 81,120 barcodes were identified in the *PKR* variant library. 86% of the barcodes were linked to select nonsynonymous variants of PKR (69,441 barcodes), 8% of the barcodes linked to either WT PKR or synonymous variants (744 and 6,092 barcodes, respectively), and 6% of the barcodes linked to off-target variants (4,843 barcodes). Off-target variants are barcodes linked to nonsynonymous variants outside the windows of interest or contain >1 nonsynonymous variant.

**Figure S9.**
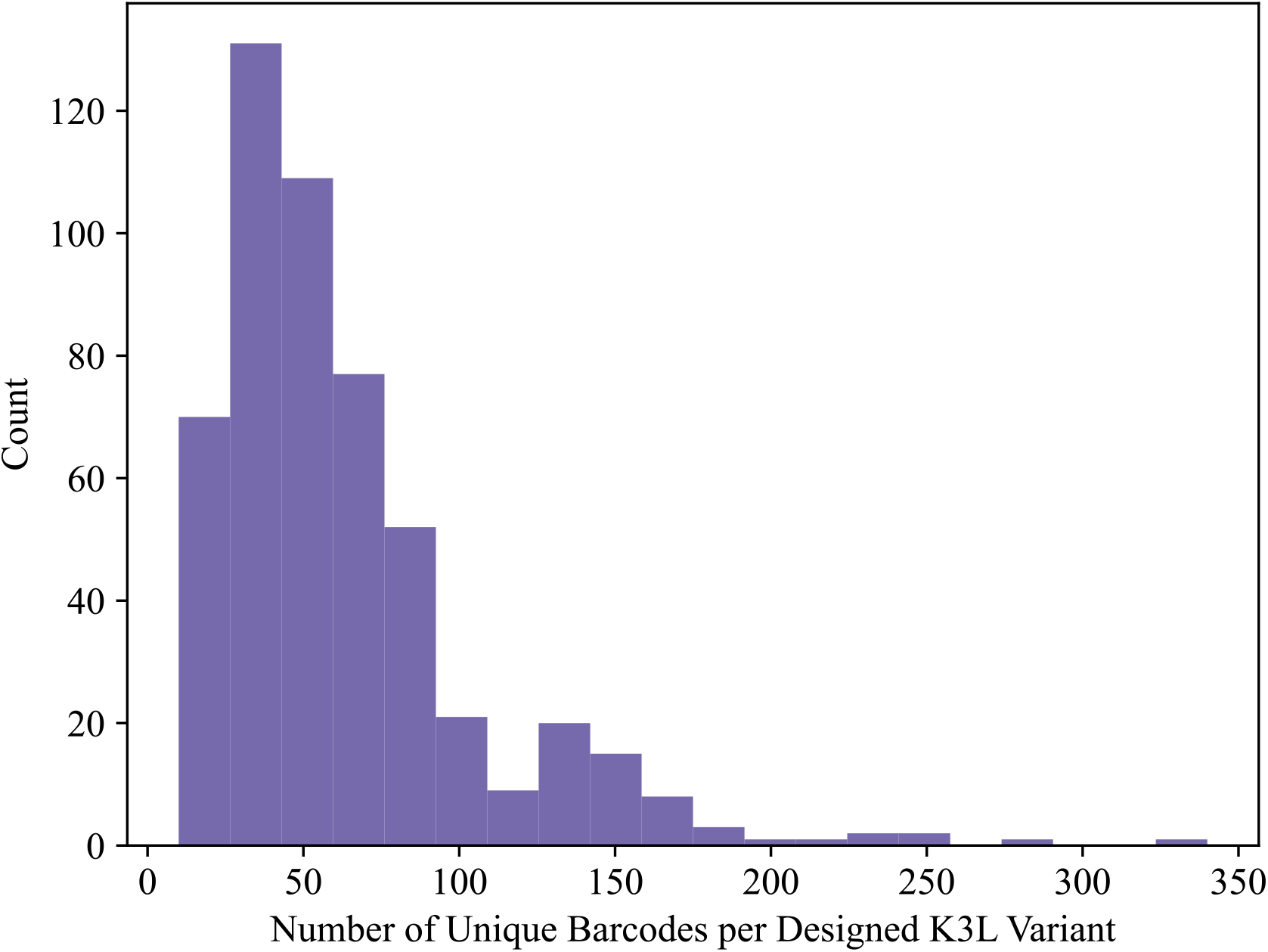
Count of unique barcodes per designed *K3L* variant. Histogram depicts the number of unique barcodes per designed *K3L* variant with a mean of 63 barcodes per *K3L* variant.

**Figure S10.**
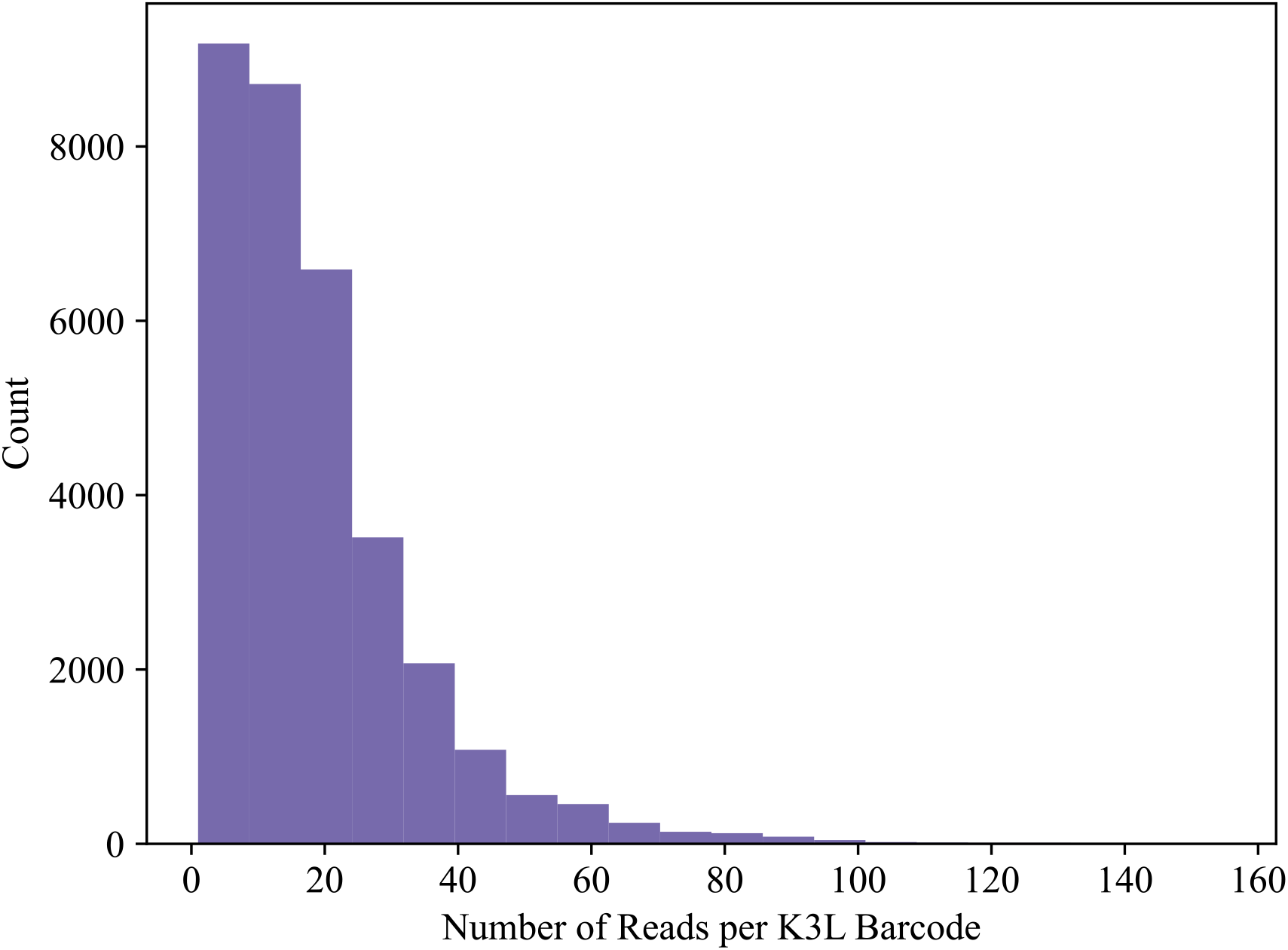
PacBio read depth per *K3L* barcode. Histogram depicts the number of reads per *K3L* barcode with a mean of 18 reads per barcode.

**Figure S11.**
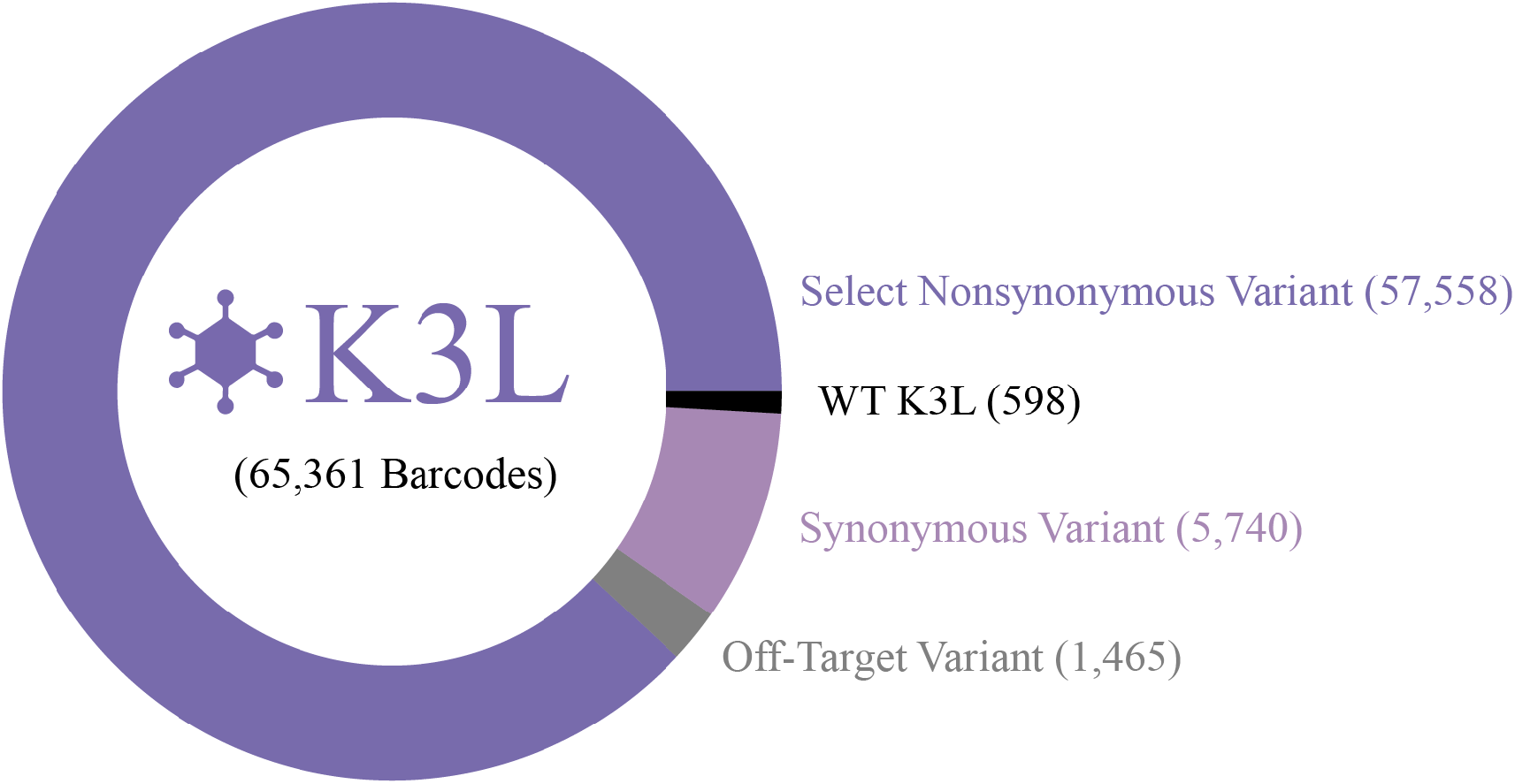
PacBio barcode distribution linked to *K3L* variant types. A total of 65,361 barcodes were identified in the *K3L* variant library. 88% of the barcodes were linked to select nonsynonymous variants of *K3L* (57,558 barcodes), 10% of the barcodes were linked to either WT K3 or synonymous variants (598 and 5740 barcodes, respectively), and 2% of the barcodes were linked to off-target variants (1,465 barcodes). Off-target variants are barcodes linked to nonsynonymous variants outside residues L2-Q88 or contain >1 nonsynonymous variant.

**Figure S12.**
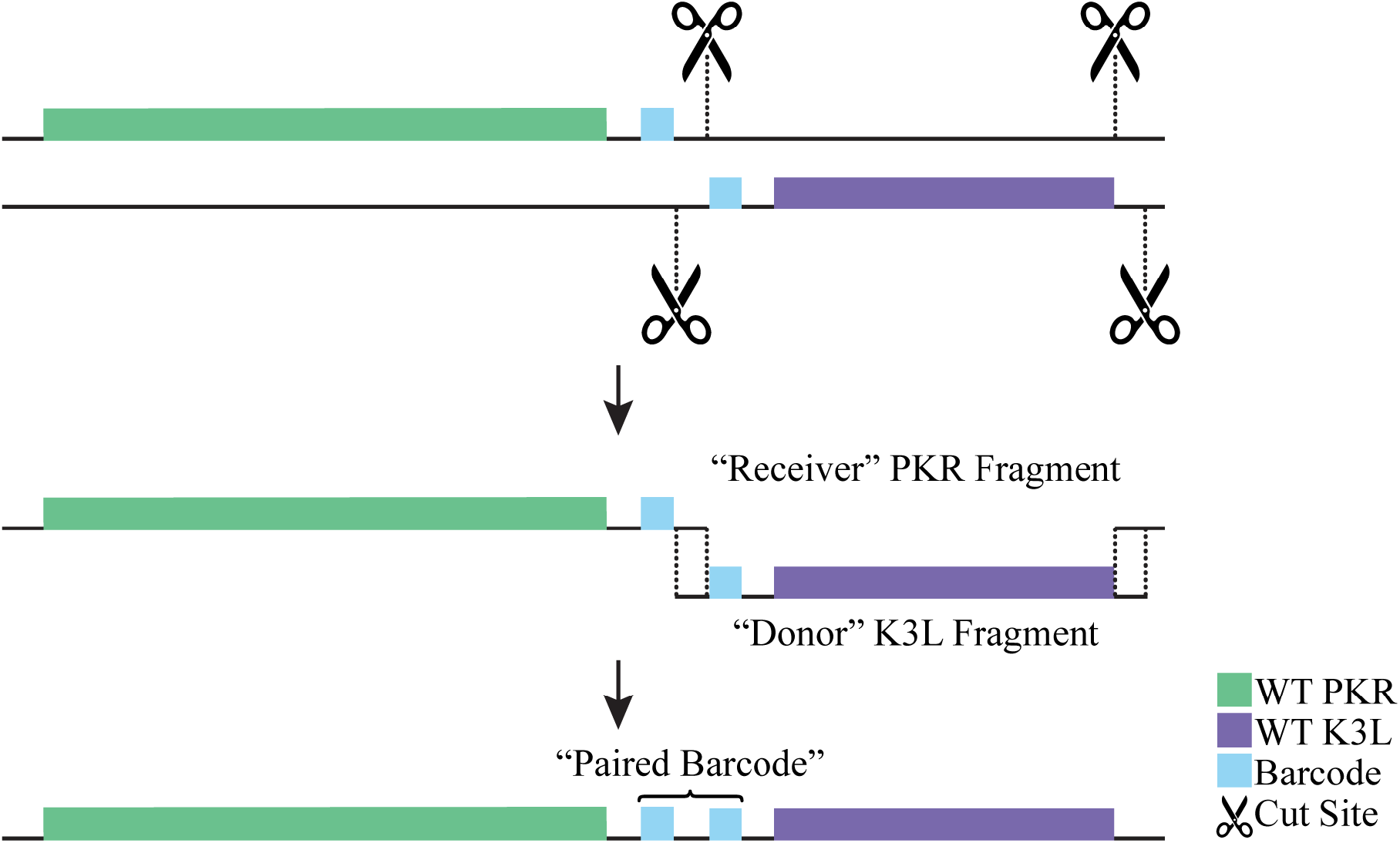
*PKR* and *K3L* variant libraries were combined to form an all-pairs library containing all 204,500 unique pairs of *PKR* and *K3L* variants with adjacent barcodes to form a “paired barcode”. A double-digest of the *PKR* variant library creates a landing pad with homology to a double-digest fragment from the *K3L* variant library using unique restriction enzyme cut sites for each variant library. The *PKR* and *K3L* fragments are flanked with 20 bp homology arms that complement one another for Gibson Assembly, forming a library of 204,500 unique pairs. *PKR* and *K3L* barcodes are placed adjacent to one-another to form a “paired barcode” that can be read via Illumina short-read sequencing.

**Figure S13.**
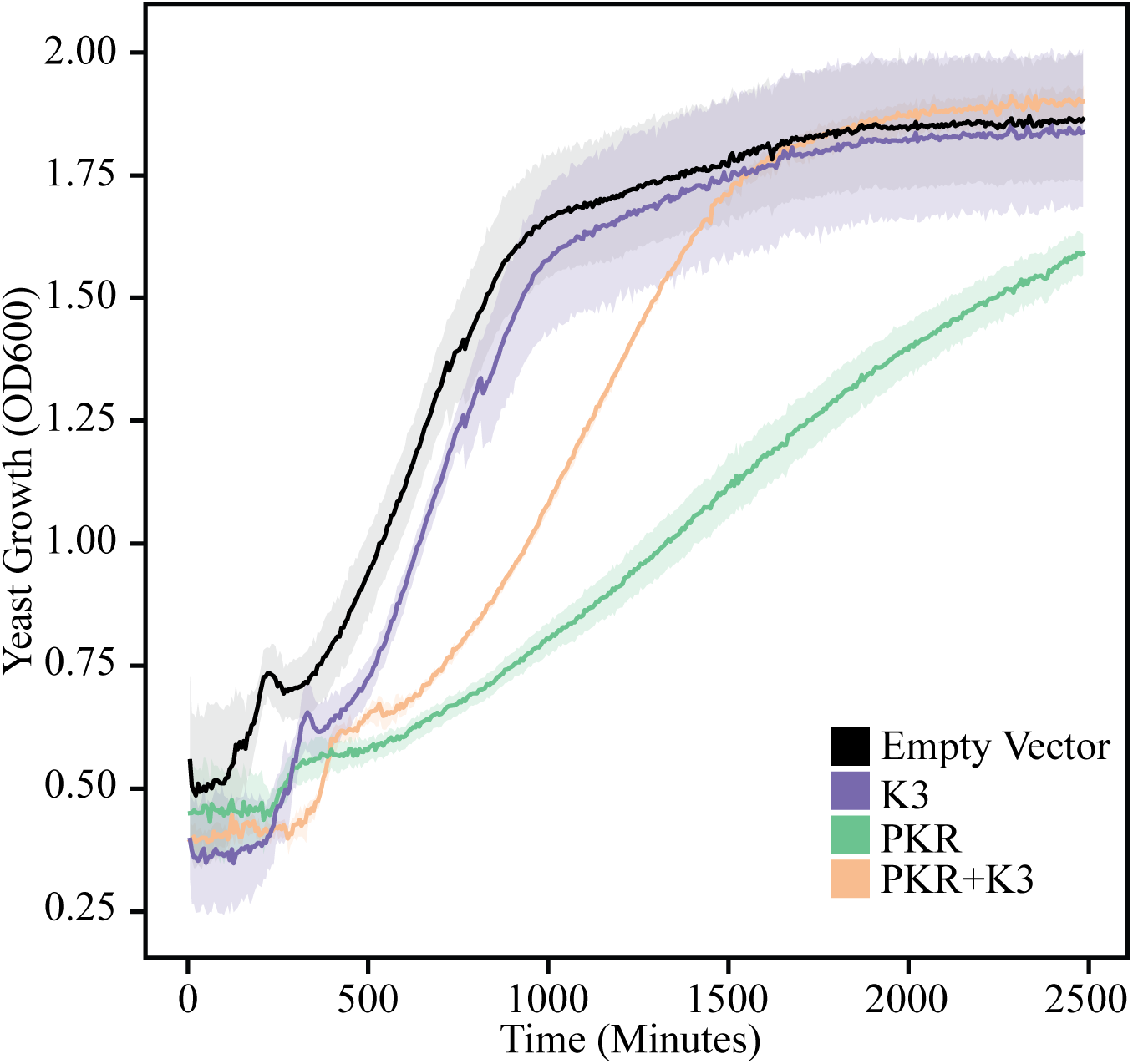
Expression of PKR and K3 alters yeast growth rates. Expression of PKR, K3, or a combination of the two in yeast alters growth over time. WT yeast growth (black) is comparable to yeast expressing K3 (purple), while yeast growth is inhibited when expressing PKR (green). However, yeast growth is appreciably recovered when both PKR and K3 are co-expressed (orange), as K3 antagonizes PKR inhibition of translation and growth.

## Supplementary tables

**Table S1.**
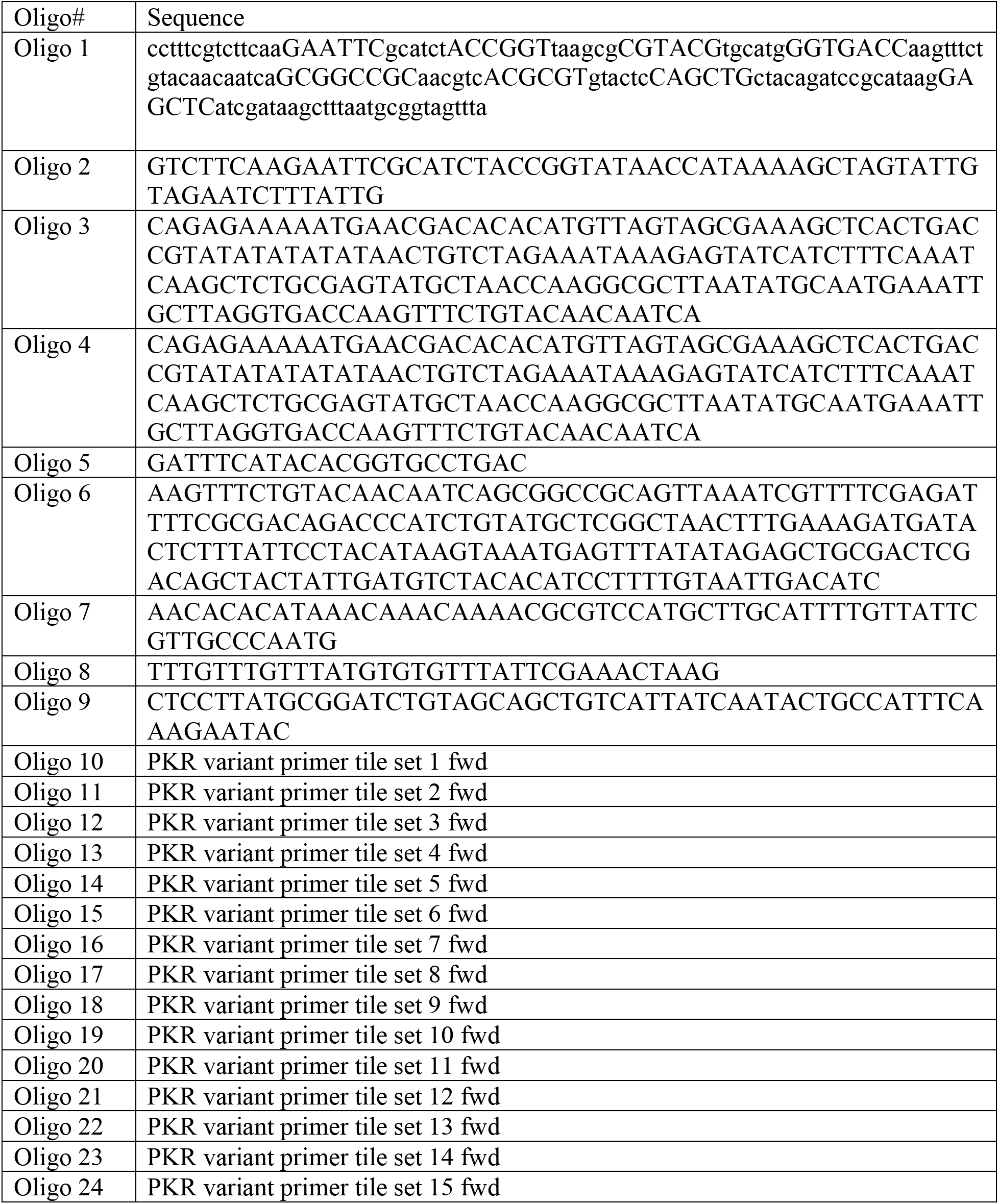
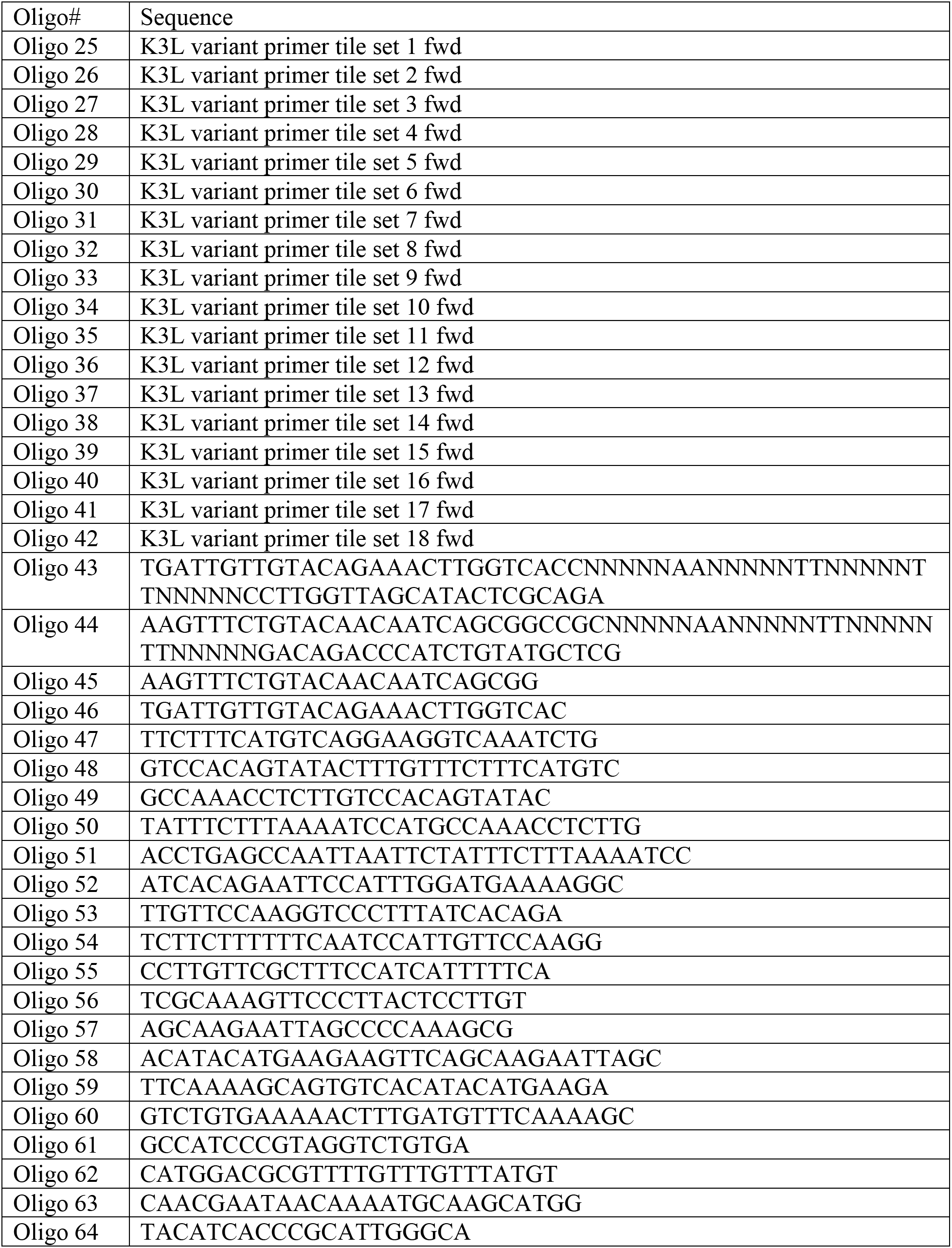
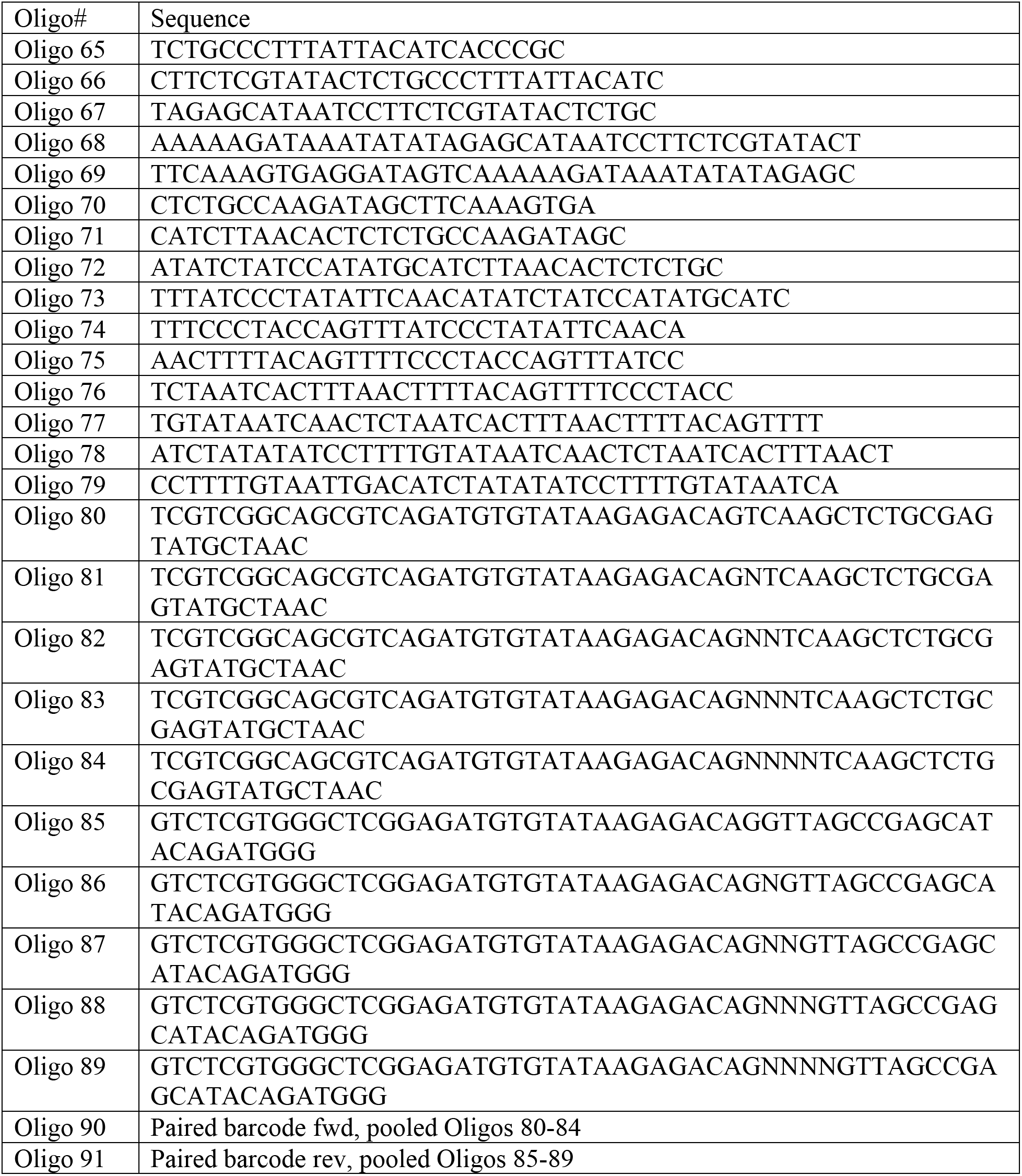
Oligos used to generate and characterize *PKR* and *K3L* variants.

